# Ventral pallidal GABAergic neurons control hedonic feeding and obesity

**DOI:** 10.64898/2026.06.18.733195

**Authors:** Justin G. Wang, Chang S. Xu, Chantelle L. Murrell, Mason R. Barrett, Gargi C. Basu, Lisa Z. Fang, Yiru Chen, Florian Schoukroun, Thomas Topilko, Johanna Perens, Jacob Hecksher-Sørensen, Meaghan C. Creed, Alexxai V. Kravitz

## Abstract

Food intake is governed by two interacting drives. The homeostatic hunger drive regulates food intake to fulfill caloric needs while the hedonic drive promotes intake of palatable foods outside of caloric need. It is unclear which neural substrates can control the hedonic drive and thereby reduce overeating of palatable foods and associated obesity. Here, we show that ventral pallidal GABAergic neurons (VP^GABA^) preferentially control hedonic feeding and are necessary for diet-induced obesity in mice. Stimulating VP^GABA^ neurons drove robust consumption of high-fat diet and liquids, but not regular laboratory chow. Despite driving intake of palatable foods, VP^GABA^ neurons are relatively insensitive to homeostatic signals – they express few hunger-hormone receptors and are not activated by ghrelin administration or fasting. Single-cell calcium imaging revealed stronger engagement of VP^GABA^ neurons during long vs short feeding bouts, suggesting control over bout duration, which has been linked to palatability. This was confirmed with closed-loop optogenetic stimulation. Finally, taCasp3-mediated ablation of VP^GABA^ neurons reduced intake of palatable liquids and blocked high-fat diet-induced obesity without impacting homeostatic feeding. Together, these findings establish VP^GABA^ neurons as a neural population that preferentially controls hedonic over homeostatic feeding and can be leveraged to block obesity in mice.

## Introduction

Food intake is governed by two interacting drives: a homeostatic drive promotes feeding in response to caloric need, while a hedonic drive promotes feeding independent of caloric need^1^. The homeostatic drive is centered in the hypothalamus and evolved to balance energy intake with expenditure. The hedonic drive involves the reward pathway and may have evolved to enable overconsumption of energy-dense foods, increasing caloric reserves to protect against future scarcity^2^. In modern food environments, however, ubiquitous access to energy-dense foods can render the hedonic drive maladaptive as it can promote overeating and obesity^3^. Reducing the hedonic feeding drive may represent an attractive therapeutic strategy for combatting obesity^4^. However, identifying neural populations that preferentially control hedonic over homeostatic feeding has proved elusive.

Here, we tested the hypothesis that GABAergic neurons in the ventral pallidum (VP^GABA^ neurons) preferentially control hedonic palatability driven feeding over homeostatic feeding driven by caloric need. We focused on the VP because of its long-standing role in food intake, and particularly in hedonic aspects of food intake^5^. The first clues about this role came from lesion studies of the VP, which caused both aphagia and weight loss^6^, though such lesions may have encroached on the overlying globus pallidus and neighboring lateral hypothalamus. VP lesions also reduced hedonic liking reactions to sucrose^6^, suggesting that the VP may modulate hedonic feeding via altering palatability. Bicuculline micro-injections to disinhibit the VP also induced robust feeding^7^, which was preferential for calorie-dense fats over proteins or carbohydrates^8^, again suggesting regulation of feeding outside of a purely homeostatic drive. Modern manipulations have started to dissect the cell-type specific role of VP neurons in feeding, specifically implicating GABAergic neurons of the VP (VP^GABA^ neurons) in controlling food intake^9,10^. For these reasons, the VP has been hypothesized as a promising novel structure to target for combatting obesity^11,12^.

However, it remains unclear if VP^GABA^ neurons preferentially control hedonic over homeostatic feeding, and if manipulation of VP^GABA^ neurons can alter the development of obesity. Here, we formally tested the role of VP^GABA^ neurons in hedonic feeding, contrasting their activity and function with that of a canonical driver of homeostatic feeding, the agouti-related peptide expressing neurons of the arcuate nucleus of the hypothalamus (Arc^AgRP^ neurons)^13,14^. With optogenetics, fiber photometry, single-cell calcium imaging, and taCasp3-mediated ablation with head-fixed behavioral tests, we demonstrate that VP^GABA^ neurons preferentially control hedonic over homeostatic feeding and are required for the development of diet-induced obesity.

## Results

### Arc^AgRP^ and VP^GABA^ neurons activate distinct feeding drives

We selectively expressed channelrhodopsin-2 (ChR2) in Arc^AgRP^ neurons (n=6, 3M/3F) or VP^GABA^ neurons (n=12, 5M/7F), to compare how activation of each population impacted feeding. Consistent with prior literature^14^, stimulation of Arc^AgRP^ neurons (bilateral 20Hz stimulation, 5mW, 10ms pulse-width) drove intake of both standard laboratory chow and palatable high-fat diet (HFD, 60% by calories, **Fig 1A-C**). Pre-stimulating Arc^AgRP^ neurons for 30 minutes prior to food availability also increased subsequent consumption of both chow and HFD (**Fig 1D**)^15^, suggesting induction of a long-lasting hunger state. VP^GABA^ neuron activation also drove food intake, but with clear differences from Arc^AgRP^ activation. First, while activation of VP^GABA^ neurons reliably drove mice to consume HFD (**Video S1)**, it only induced chow consumption in 3 of 12 mice (**Fig 1E-G**). Pre-stimulation of VP^GABA^ neurons for 30 minutes also did not enhance subsequent food intake, demonstrating that activation of VP^GABA^ neurons did not drive a hunger state that persisted beyond the length of stimulation (**Fig 1H**). Activation of VP^GABA^ neurons also induced non-consumption interactions with chow (**Video S2**) and a plastic Lego brick (**Video S3**) in some mice, but this was not reliably observed across all mice (**Fig S1A, B**). Finally, activation of Arc^AgRP^ neurons has been linked to an anxiolytic effect, as evidenced by heightened exploration^16^. In contrast, activation of VP^GABA^ neurons caused mice to spend less time in the center of an open field, without any change in average speed (**Fig S1C, D**). While we were not powered to detect sex differences, the main findings were present in both males and females (**Supplemental Table 1**). Overall, our experiments demonstrate that both Arc^AgRP^ and VP^GABA^ activation drives feeding, but with important differences. While Arc^AgRP^ activation drives behavior consistent with a homeostatic feeding drive, activation of VP^GABA^ neurons drives behavior consistent with a hedonic feeding drive.

**Figure 1.**
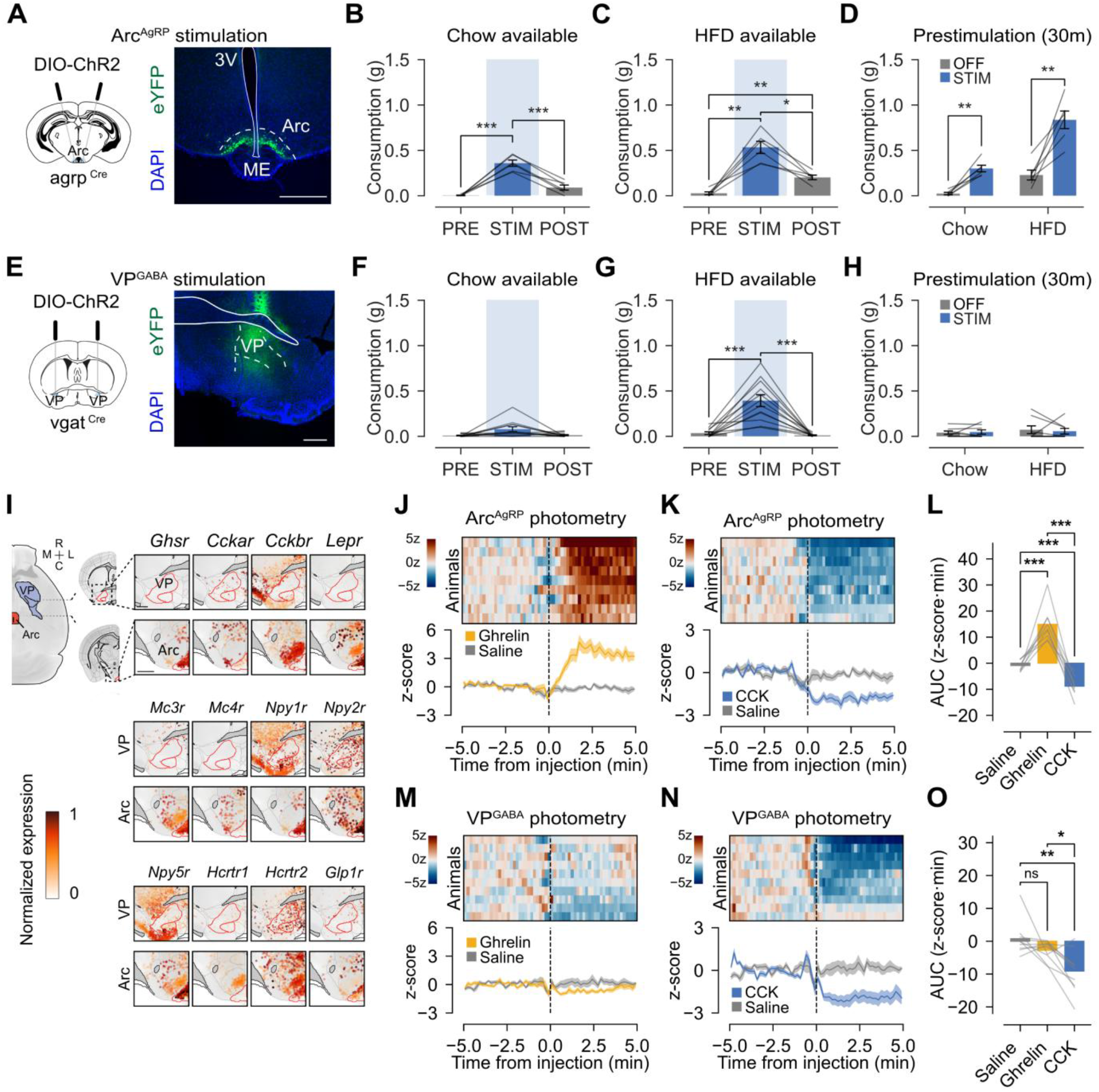
Arc^AgRP^ and VP^GABA^ neurons activate distinct feeding drives. **A**. Schematic and representative histology image for ChR2 expression in Arc^AgRP^ neurons. **B**. Optical stimulation of Arc^AgRP^ neurons drives chow consumption in sated mice. **C**. Optical stimulation of Arc^AgRP^ neurons drives HFD consumption in sated mice, with persistent feeding after stimulation. **D**. Pre-stimulation of Arc^AgRP^ neurons for 30 min before diet availability drives subsequent chow and HFD consumption. **E**. Schematic and representative histology for ChR2 expression in VP^GABA^ neurons. **F**. Optical stimulation of VP^GABA^ neurons does not significantly alter chow intake in sated mice. **G**. Optical stimulation of VP^GABA^ neurons selectively drives HFD consumption in sated mice. **H**. Pre-stimulation of VP^GABA^ neurons for 30 min before diet availability does not drive subsequent chow or HFD consumption. **I**. Spatial projection of receptor-expression maps for 12 feeding related receptors projected onto VP and Arc regions. **J–K**. Fiber photometry recordings from Arc^AgRP^ neurons showing responses to ghrelin and CCK injections, respectively. Heatmaps show individual animals and traces below show average z-scored fluorescence responses, with paired saline controls. **L**. Quantification of post-injection area under the curve (AUC) from Arc^AgRP^ recordings. **M–O**. Same format as **J-L** but showing recordings from VP^GABA^ neurons. Data are presented as mean ± SEM. Displayed symbols indicate significance of Bonferroni- or Holm-corrected post hoc comparisons (\**p*<0.05, \*\**p*<0.01, \*\*\**p*<0.001). Detailed statistical analyses in **Supplemental Table 1**.

To further examine differences in how the Arc and VP control feeding, we integrated previously published spatial transcriptomic data^17^ with single cell RNA sequencing (scSeq) data^18^ to estimate expression levels of twelve feeding-related receptor transcripts (*Ghsr, CCKAr, CCKBr, LepR, MC3R, MC4R, Npy1r, Npy2r, Npy5r, Hcrtr1, Hcrtr2*, and *Glp1r*) in the VP and Arc. For each spatial cell, the displayed expression value corresponds to the mean log2-normalized expression of the selected gene in the cell’s assigned transcriptomic cluster, derived from the scSeq dataset. Thus, the maps represent reference-based, cluster-level expected expression projected onto spatially registered MERFISH cells, rather than direct measurement of the displayed genes in individual MERFISH cells. While the Arc expressed high levels of nearly all of these receptors, the VP expressed relatively low levels (**Fig 1I**). Notable exceptions include *Cckar, Npy2r*, and *Hcrtr2*, which were expressed at moderate levels in the VP. We confirmed this in two additional datasets including a different scSeq dataset from the Arc^19^ and a single nucleus RNA sequencing (snSeq) of the VP^20^ (**Fig S2A, B**). Unlike Arc^AgRP^ neurons, VP^GABA^ neurons are relatively insensitive to hunger and satiety states, at least via direct receptor activation.

We considered that VP^GABA^ neurons might *indirectly* sense hunger or satiety states via afferent inputs, regardless of whether they expressed the above receptors. To test this possibility, we used fiber photometry to measure calcium dynamics in mice expressing GCaMP8f in Arc^AgRP^ (n=9, 5M/4F) or VP^GABA^ neurons (n=10, 3M/7F), in response to an injection of ghrelin (1mg/kg subcutaneous, SC) in sated mice to rapidly induce hunger, cholecystokinin (CCK octapeptide, SC, 10 μg/kg) in fasted mice to rapidly induce satiety, or saline as a control. Consistent with prior reports^21,22^, ghrelin caused a rapid increase in calcium activity in Arc^AgRP^ neurons, while CCK caused a rapid decrease (**Fig 1J-L**). However, ghrelin did not alter calcium activity of VP^GABA^ neurons, demonstrating that VP^GABA^ neurons are insensitive to this induced hunger state (**Fig 1M**). In contrast, CCK did inhibit VP^GABA^ neurons, suggesting that they can detect the CCK-induced satiety state (**Fig 1N,O**). We also offered ghrelin and saline injected mice grain pellets to quantify their calcium responses to feeding. Arc^AgRP^ neurons were rapidly inhibited at the first pellet retrieval and stayed inhibited for the remainder of the feeding period (**Fig S3**). VP^GABA^ neurons were activated during each pellet retrieval but these responses completely recovered between each pellet.

We next tested if VP^GABA^ neurons were sensitive to natural hunger, using cFos expression as a readout. We fasted mice for 12 hours to induce hunger, while maintaining a control group on ad lib chow without fasting (n=8 per group, all male). Brains were removed, cleared, and stained for cFos expression with resulting images registered to the Gubra Multimodal 3D Brain Atlas Framework^23^ to quantify cFos nuclei in ∼800 brain areas (**Fig S2C-F**). We did not detect any induction of cFos in the VP after fasting but observed that fasting induced cFos in many other brain areas (**Fig S2E, Supplemental Table 2**). Of the 13 described inputs to the VP^24^, 4 showed evidence of cFos activation in response to fasting (**Fig S2F**), including areas that have previously been implicated in food intake such as the paraventricular nucleus of the thalamus (p=0.009), lateral part of the central amygdala nucleus (p=0.02) and the dentate gyrus (p=0.005) and CA1 region of the hippocampus (p=0.04). In this way, the VP may receive information about hunger states and use this information to gate or influence other signals, even though hunger does not alter VP^GABA^ activity as observed by population calcium or cFos induction.

### VP^GABA^ neurons bi-directionally control reward consumption

We used a head-fixed preparation to test how VP^GABA^ optogenetic activation or inhibition would impact the microstructure of consumption. Animals expressed ChR2 (n=8-15, mixed sex) or soma-targeted chloride-conducting channelrhodopsin (stGtACR2, n=6 males) to provide activation or inhibition of VP^GABA^ neurons. Consistent with prior literature^25^, VP^GABA^ activation was preferred in a real-time place preference (RTPP) task, while optogenetic inhibition was aversive (**Fig S4**). Animals in both groups were trained to lick a spout to consume a high-calorie chocolate-flavored liquid (80% Boost; hereafter “Boost”) while we delivered optogenetic activation or inhibition (**Fig 2A, B**). ChR2 expressing animals were first given the opportunity to consume Boost in 100-second blocks, with five interleaved 5-second bursts of optogenetic activation every 20s (20 Hz, 5 mW, 10 ms pulse-width). VP^GABA^ activation rapidly increased consumption relative to matched control trials without activation, driven by a significant increase in both bout number and duration (**Fig 2C**). To assess the upper limit of VP^GABA^ activation on licking behavior, we delivered 95-seconds of continuous stimulation (20Hz, 5mW, 10 ms pulse-width) to the same mice in a separate session, finding that consumption remained elevated for the full stimulation duration, again driven by increases in both bout number and duration (**Fig 2D, Video S4**). We next tested the effect of optogenetic inhibition in stGtACR2 expressing mice. Inhibition was delivered in 2-minute blocks interleaved with 2 minute-blocks without stimulation over 30 minutes. Inhibition caused mild reductions in total licks, driven by a decrease in the number of bouts and duration (**Fig 2E)**. The bi-directional actions on bout number reflect motivation or “wanting” for the outcome^26^, while the changes in bout duration are consistent with changes in palatability^27,28^. Through this lens, we concluded that VP^GABA^ activation increased palatability, and next tested if it could drive intake of aversive outcomes such as 1% quinine, a bitter solution. VP^GABA^ activation increased the number and duration of bouts of quinine as well, demonstrating that it could in fact increase the palatability of quinine (**Fig S5A**). However, the increase in licking for quinine was more modest than for Boost and not sustained across the stimulation window (**Video S4**). A direct comparison of licking microstructure for Boost, quinine, and an empty spout revealed similar numbers of bouts for each outcome, but an ordered relationship such that Boost induced longer duration licking than the empty spout, which induced longer duration licking than quinine, consistent with their perceived palatability (**Fig S5C**).

**Figure 2.**
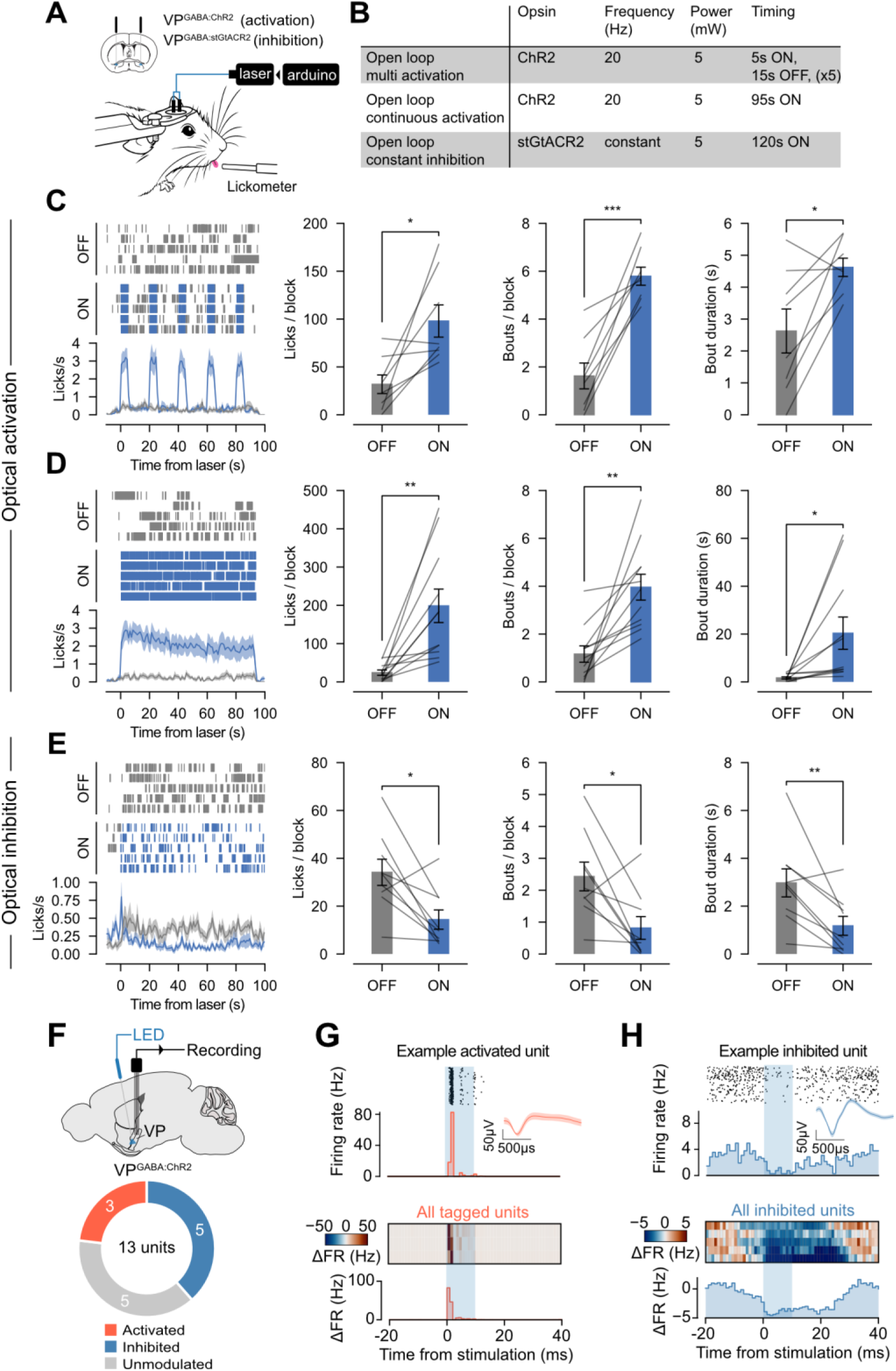
VP^GABA^ neurons bi-directionally control reward consumption. **A**. Schematic of open-loop optogenetic manipulation of VP^GABA^ neurons during head-fixed licking with concurrent optogenetics. **B**. Stimulation paradigms used for multi-block activation, continuous activation, and constant inhibition experiments. **C**. Example licks in rasters, and average licks/second across 5s of 20Hz ChR2 mediated activation, interleaved with 15s OFF periods. Licks/block, bouts/block, and bout duration quantified between laser OFF vs ON blocks. **D**. Same presentation as **C** but for 95 seconds of 20Hz ChR2 stimulation. **E**. Same presentation as **C** but for 120 seconds of constant stGtACR2-mediated inhibition. **F**. Schematic of extracellular optrode recordings from VP^GABA^ mice, with classification of recorded VP units based on their response to optical stimulation. **G**. Example activated unit (top) with average response of 3 activated units (bottom). **H**. Example inhibited unit (top) with average response of 5 inhibited units (bottom). Data are presented as mean ± SEM. Asterisks indicate significance of paired comparisons (\**p*<0.05, \*\**p*<0.01, \*\*\**p*<0.001). Detailed statistical analyses in **Supplemental Table 1**.

To determine how optogenetic activation impacted the firing of VP^GABA^ neurons, we performed in vivo extracellular recordings, obtaining recordings from 13 single units from 3 mice that expressed ChR2 in VP^GABA^ neurons. We delivered 10ms, 5mW test pulses at 20Hz, as in the behavioral tests above. We observed rapid (< 3ms) activation in 3 of the 13 single units, consistent with ChR2-expressing VP^GABA^ neurons (**Fig 2F-H**). We also observed inhibition in 5 of 13 single units, likely caused by GABA release from stimulated VP^GABA^ neurons. This inhibition lasted for 20-30ms after the stimulation ended, which may reflect the time-course of GABA clearance. These recordings demonstrate direct activation of VP^GABA^ units by optogenetic activation, but also reveal inhibition of neighboring VP neurons, as expected from stimulating a GABAergic population.

### Single-photon endoscopic imaging reveals functionally heterogeneous VP^GABA^ dynamics that encode and track consummatory licking

Optogenetic activation of VP^GABA^ neurons increases both bout number and duration (**Fig 2**). We therefore hypothesized that VP^GABA^ neurons might reflect the initiation or duration of licking bouts in their endogenous activity. To test this, we expressed a Cre-dependent calcium indicator GCaMP7s in VP^GABA^ neurons and imaged somatic calcium activity of 339 single VP^GABA^ neurons (n=8, 7M/1F, 13 sessions) through an endoscopic Gradient Refractive Index (GRIN) lens coupled to an imaging fiber bundle (Mightex OASIS system, **Fig 3A, Video S5**), while head-fixed mice freely licked for Boost. Consistent with prior in vivo recordings of the VP^29–31^, response profiles of individual neurons were heterogeneous, with ∼38% of the recorded neurons activated during licking, 50% inhibited, and 12% unmodulated (**Fig 3B, C**). We tested whether VP^GABA^ calcium activity represented bout length by classifying licking bouts as short (bottom quartile) or long (top quartile, **Fig 3D**). Calcium responses scaled monotonically in amplitude, such that single isolated licks evoked the smallest calcium responses, and long bouts produced the largest (**Fig 3E**). The long bouts were represented by higher amplitude responses in both activated and inhibited neurons (**Fig 3E, F**). In addition, the proportion of modulated neurons was higher in long bouts (**Fig 3F**).

**Figure 3.**
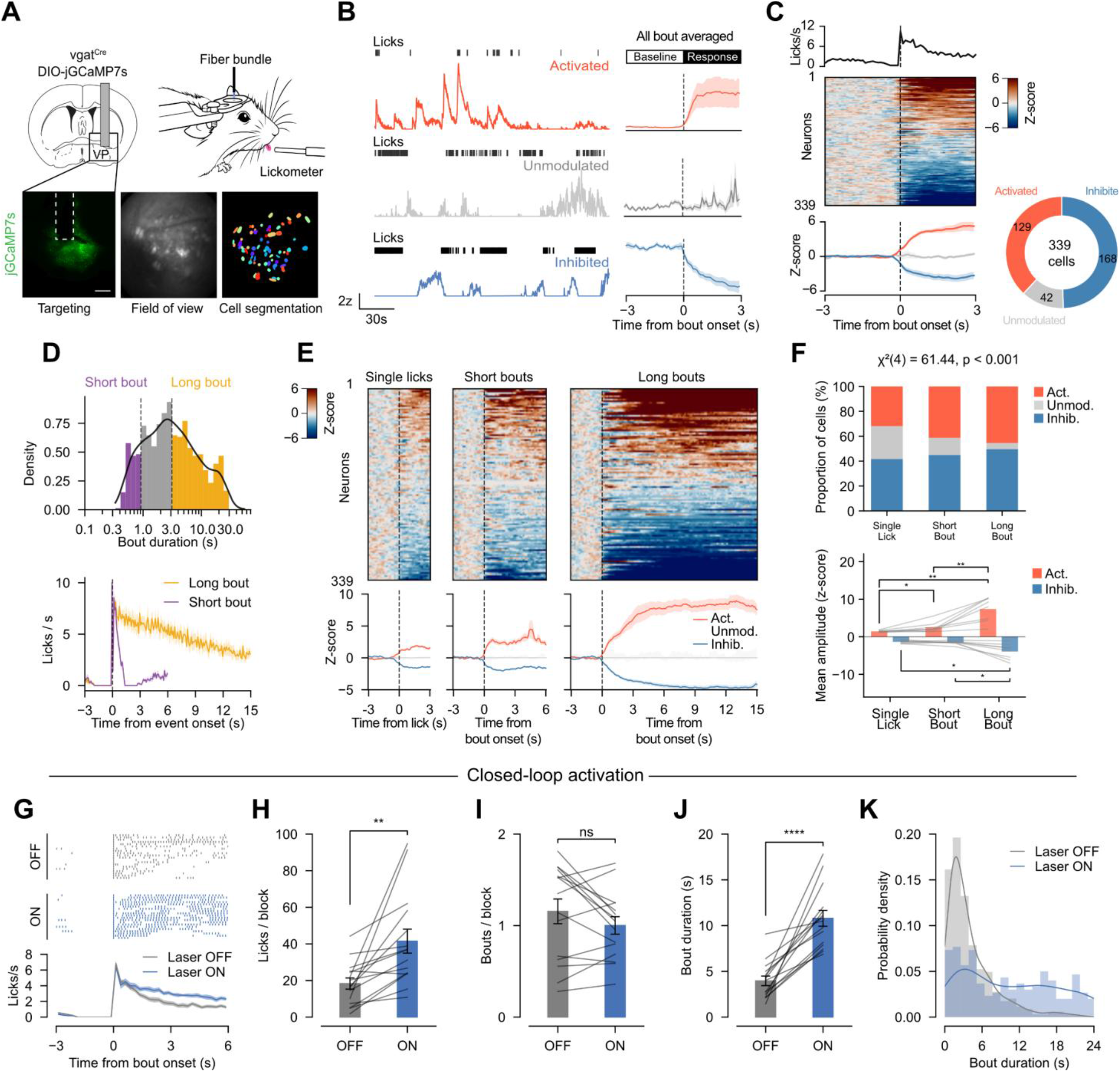
Single-photon endoscopic imaging reveals functionally heterogeneous VP^GABA^ dynamics that encode and track consummatory licking. **A**. Schematic of 1-photon calcium imaging of VP^GABA^ neurons. Representative lens targeting, imaging field of view, and cell segmentation map for one mouse. **B**. Representative calcium traces from example activated, unmodulated, and inhibited VP^GABA^ neurons aligned to licking behavior, with corresponding bout-triggered averages. **C**. Population responses aligned to the start of lick bouts, including lick rate (top), single-cell z-score heatmap (middle), and class-averaged activity traces (bottom). Cells were classified as activated, inhibited, or unmodulated (339 total neurons). **D**. Distribution of bout durations showing definition and lick-rate dynamics for short vs. long bouts. **E**. Single-neuron heatmaps and class-averaged responses for single licks, short bouts, and long bouts. **F**. Cell-type proportions and response amplitudes across lick types, showing proportion and mean amplitude of response for activated and inhibited populations by bout structure. **H**. Representative lick rasters and average licking rate for closed-loop optogenetic activation of VP^GABA^ neurons. **H-J**. Effects of VP^GABA^ activation on licks/block, bouts/block, and bout duration. **K**. Probability density distribution of bout durations during closed-loop ChR2 stimulation, vs. interleaved non-stim periods. Data are presented as mean ± SEM. Significance assessed using χ^2^ tests, permutation tests, Holm-corrected post hoc tests, and paired t-tests as appropriate (\**p*<0.05, \*\**p*<0.01, \*\*\**p*<0.001, ****p<0.0001). Detailed statistical results in **Supplemental Table 1**. analysis confirms that increasing the activity of VP^GABA^ neurons causally controls bout duration. In this way, VP^GABA^ neurons can control palatability.

The variability in responses of VP^GABA^ neurons during consumption stands in contrast to Arc^AgRP^ neurons, which are uniformly inhibited during consumption^32,33^, suggesting that VP^GABA^ neurons use a higher dimensional population code to represent feeding duration. We used principal component analysis (PCA) to investigate this, projecting the responses of all neurons around all licking bouts onto two dimensions (**Fig S6A**), revealing that short vs. long bouts traversed divergent trajectories, suggesting that modulated neurons in short bouts were a distinct subpopulation from those modulated during long bouts (**Fig S6B-D**). To test if short vs. long bouts could be decoded from the activity of all recorded neurons, we trained a linear logistic decoder on increasing time windows after the onset of each bout (**Fig S6E-H**). The decoder could not decode short vs. long bouts until the behavior diverged (∼1 second after the bout onset), demonstrating that they encode the duration of licking bouts, but not necessarily the decision to terminate the bout before it ends. We were also not able to decode the timing of bout initiation before it began, suggesting that VP^GABA^ neurons most strongly track the duration of licking, consistent with a palatability signal.

We hypothesized that if VP^GABA^ neurons control bout duration, activating them during licking should extend bouts, creating more long bouts of licking. We activated VP^GABA^ neurons in a closed-loop design, where each self-initiated lick resulted in 1s of optogenetic stimulation (20Hz, 5mW, 10ms pulse-width, **Fig 3G**). Relative to interleaved periods with no stimulation, closed-loop activation of VP^GABA^ neurons resulted in a >2-fold increase in bout duration, without impacting bout number (**Fig 3H-K**). As suggested by the calcium imaging, this

### VP^GABA^ neurons are required for the development of diet-induced obesity

Optical inhibition reduced intake of Boost (**Fig 2E**), but the optogenetic approach is not suitable for multi-day manipulations to investigate changes in body weight. We therefore expressed Cre-dependent taCasp3, or mCherry fluorophore as a control, in VGAT-Cre mice. RNAscope *in situ* hybridization confirmed an ∼80% reduction in VP^GABA^ neurons, with a complete sparing of glutamatergic neurons in the VP (n=5, 3M/2F per group, **Fig 4A, B**). In a separate cohort (n=5 mCherry and 7 taCasp3), we tracked body weights for 12 weeks on chow following VP^GABA^ ablation surgery noting no weight loss (**Fig S7A-C**), as had been previously reported following excitotoxic lesions of the VP in rats^6^. To test how VP^GABA^ ablation impacted licking microstructure, we trained a new cohort of taCasp3 (n=11, 6M/5F) vs mCherry expressing control mice (n=8, 2M/6F) on a head-fixed self-paced drinking task for Boost. taCasp3-expressing mice had significantly fewer licks, relative to controls (**Fig 4C**), which was driven primarily by a decrease in bout number rather than bout duration (**Fig 4D, E**) and was associated with slower modal lick frequency (**Fig 4F**). Overall, this pattern differed from our acute optogenetic stimulation and suggests that ablation of VP^GABA^ neurons decreases motivation for Boost and may impair the motor generation of the licking pattern, without impacting bout duration. Given that VP^GABA^ ablation did not decrease bout duration, we next tested if HFD preference would also be preserved in mice following VP^GABA^ ablation. Ablated mice and controls both strongly preferred HFD over chow, indicating that taste discrimination and palatability of HFD were not eliminated by VP^GABA^ ablation (**Fig 4G**). This experiment also argues against the ablation causing an anhedonic phenotype. However, taCasp3 expressing mice consumed significantly less HFD than controls when sated (**Fig 4G**). In contrast, when fasted, both taCasp3 expressing mice and controls ate similar levels of HFD, suggesting that homeostatic need remains intact in the taCasp3 expressing mice. To more formally test if homeostatic need was intact, we subjected mice to alternating 24-hour periods of fasting and *ad libitum* grain-pellet access for three consecutive cycles. Both groups lost weight during fasting periods and exhibited compensatory overeating and weight recovery during *ad libitum* pellet access, demonstrating intact sensing of caloric deficit and appropriate homeostatic regulation of feeding (**Fig S7D-F**). TaCasp3 and mCherry expressing mice also had similar circadian feeding patterns (**Fig S8A, B**), although we noted a mild decrease in total HFD consumed in taCasp3 expressing mice (**Fig S8D**).

**Figure 4.**
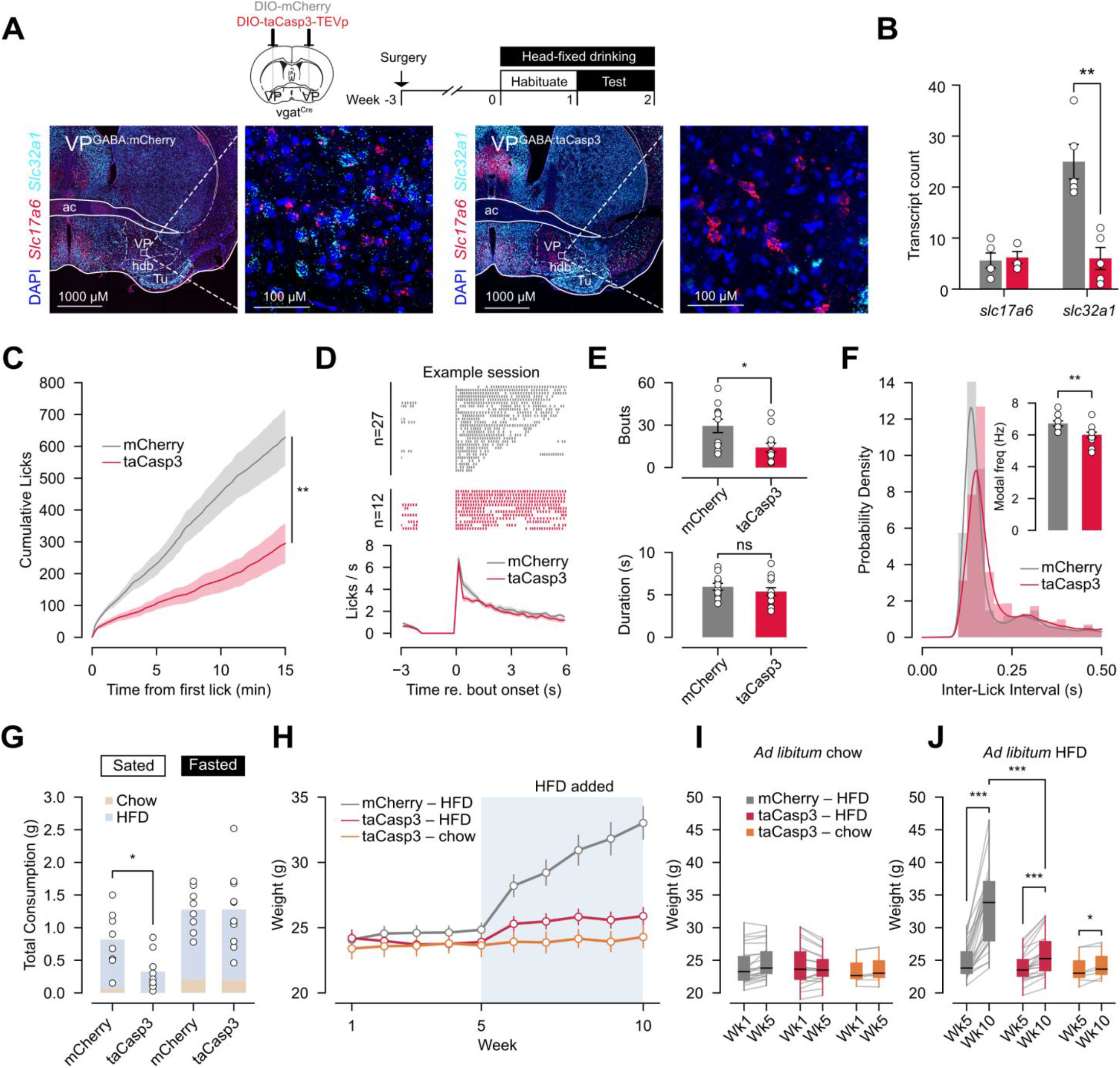
VP^GABA^ neurons are required for the development of diet-induced obesity. **A**. Schematic of taCasp3 mediated ablation of VP^GABA^ neurons, vs. mCherry controls, with representative RNAscope in situ hybridization of VP showing Slc17a6 (vGlut2), Slc32a1 (vGAT), and DAPI labeling. **B**. Quantification of Slc17a6 and Slc32a1 in the VP of taCasp3 vs. mCherry expressing mice. **C**. Cumulative licking for Boost during a 60 minute head-fixed drinking period in taCasp3 vs. mCherry expressing mice. **D**. Example lick-bout rasters from a single session (top) with mouse-averaged licking rate (bottom) in taCasp3 vs. mCherry expressing mice. **E**. Total bouts and mean bout duration between taCasp3 vs. mCherry expressing mice. **F**. Inter-lick interval distributions and modal lick frequency in taCasp3 vs. mCherry expressing mice. **G**. Total chow and HFD intake during sated and fasted states in taCasp3 vs. mCherry expressing mice. **H**. Body-weight trajectories over 10 weeks for mCherry expressing mice on HFD, and taCasp3 expressing mice on Chow or HFD. HFD introduced at week 5. **I–J**. Longitudinal body-weight comparisons across week 1–5 and week 5–10 in mCherry-HFD, taCasp3-HFD, and taCasp3-Chow groups. Data are presented as mean ± SEM. Statistical significance was assessed using t-tests, mixed-design RM-ANOVA, Mann–Whitney U tests, and permutation tests as appropriate (\**p*<0.05, \*\**p*<0.01, \*\*\**p*<0.001). Detailed statistical outcomes are provided in **Supplemental Table 1**.

We next tested a range of behavior in taCasp3 vs. mCherry expressing mice, to test if the ablation of VP^GABA^ neurons caused nonspecific deficits beyond the reduction in HFD intake. TaCasp3 and mCherry expressing mice performed comparably in a home-cage closed-economy progressive ratio task measuring motivational drive and persistence for grain pellets (**Fig S9A-C**) and had mild learning deficits in a home-cage closed-economy two-armed bandit task for these same pellets (**Fig S9D-K**). Finally, taCasp3 expressing mice did not exhibit deficits in a home-cage social operant task (**Fig S10**). Collectively, these results indicate that VP^GABA^ ablation does not produce gross impairments in homeostatic feeding, learning, motivation, or social behavior.

Having established that VP^GABA^ ablation does not produce deleterious effects on general behavior or homeostatic feeding, we finally tested whether taCasp3 expressing mice would be resistant to HFD-induced obesity. A new cohort of mice (n=22 taCasp3, n=23 mCherry) was maintained on standard chow for 5 weeks following stereotaxic surgery, then challenged with ad libitum HFD access for 5 weeks. TaCasp3 expressing mice were nearly completely protected from weight gain despite consuming HFD as their sole caloric source for the 5-week period (**Fig 4H-J, Video S6, S7)**. This resistance to obesity was present in both males and females, although mCherry expressing males gained more weight than females (**Supplemental Table 1**). Despite their resistance to obesity, taCasp3 expressing mice did become hyperglycemic after 5 weeks of consuming HFD as their only food source, demonstrating that the deleterious metabolic effects of HFD can occur even without obesity (**Fig S7G, H**). We conclude that VP^GABA^ neurons preferentially drive hedonic over homeostatic feeding, and that ablating them prevents HFD-induced obesity in mice.

## Discussion

Here, we report that VP^GABA^ neurons preferentially control hedonic over homeostatic feeding in mice. Through multiple assays, we reveal that activation of VP^GABA^ neurons drives intake of palatable, but not unpalatable food, while inhibition of VP^GABA^ neurons decreases intake and blocks HFD-induced weight gain. Analysis of licking microstructure was particularly revealing, showing that activation of VP^GABA^ neurons increases both motivation for feeding (as evidenced by increased bout number), as well as palatability (increased bout duration). However, the endogenous activity of VP^GABA^ neurons most strongly scaled with bout duration and did not predict bout initiation, suggesting that VP^GABA^ activity more closely tracks ongoing consumption and bout duration than it drives the decision to eat. Finally, taCasp3 mediated ablation of VP^GABA^ neurons reduced over-eating and blocked HFD-induced weight gain, while sparing homeostatic control of feeding.

A candidate mechanism by which VP^GABA^ neurons promote hedonic feeding may be through dopamine-dependent reinforcement. VP^GABA^ neurons project directly to the VTA and substantia nigra pars compacta, where their activation disinhibits dopamine neurons and increases dopamine release in the nucleus accumbens^34^. Consistent with this possibility, VTA-projecting VP neurons are activated during reward consumption and support food reinforcement^35^, while dopamine release persists throughout feeding bouts and can overcome physiological satiety signals^36^. Such a mechanism could explain our observations that VP^GABA^ activation increases bout numbers and duration by enhancing the reinforcing value of ongoing consumption and food-associated actions^37–39^.

Although we manipulated VP^GABA^ neurons in this study, this population is molecularly heterogeneous^20,40^ and projects to at least four main downstream targets including the VTA^41^. Future studies will be required to determine which molecularly or projection-defined VP^GABA^ subpopulations mediate the behavioral effects we observed. GABAergic neurons in ventral basal forebrain regions, including the substantia innominata and magnocellular preoptic area, can drive intake of standard chow as well as palatable food^42^. Our targeting and post-hoc histology are centered on the caudal dorsolateral VP, an area previously linked to hedonic facial reactions^43^, which may account for the selective intake of palatable food we observed during optogenetic activation. Future studies will be needed to identify the specific regional, projection, and molecular targets in the VP that most strongly impact feeding and obesity.

More broadly, our results demonstrate that forebrain circuits control hedonic feeding and can be targeted to block the development of obesity. In addition, these findings demonstrate that hedonic and homeostatic feeding can be dissociated in the VP. As most pharmacological strategies for combatting obesity currently target homeostatic regulation, VP circuits may offer a complementary point of intervention that selectively reduces consumption of palatable foods while preserving homeostatic mechanisms that defend body weight.

## Video Captions

**Supplemental Video S1:** Example VP^GABA:ChR2^ stimulation for HFD.

**Supplemental Video S2:** Example VP^GABA:ChR2^ stimulation for chow and Lego brick.

**Supplemental Video S3:** Example VP^GABA:ChR2^ stimulation for Lego brick.

**Supplemental Video S4:** Example VP^GABA:ChR2^ stimulation for quinine, empty spout, and Boost reward.

**Supplemental Video S5:** Example VP^GABA:jGCaMP7s^ one photon calcium imaging movie.

**Supplemental Video S6:** Example VP^GABA:mCherry^ mouse after 5 weeks of HFD exposure.

**Supplemental Video S7:** Example VP^GABA:taCasp3^ mouse after 5 weeks of HFD exposure.

## Supplemental Table Captions

**Supplemental Table 1**. Full statistics and animal numbers for Figure 1-4 and S1-10.

**Supplemental Table 2**. Whole brain cFos counts and statistics, segmented into the Gubra Multimodal 3D Brain Atlas Framework^23^.

## Concise Methods (see Supplemental Methods for full methods)

### Mice

Vgat-IRES-Cre (Jax 028862) and Agrp-IRES-Cre (Jax 012899) mice were crossed to C57BL/6J. Both sexes (≥8 weeks at surgery, ∼12 weeks at testing) were used and pooled; cohorts were not powered to test sex as a factor. Procedures were approved by the Washington University IACUC.

### Stereotaxic surgery and viruses

Under isoflurane, AAVs were injected into VP (AP +0.1, ML ±1.5, DV ™4.95 from bregma) at 10 nL s^−1^ (250–300 nL, 6–8 × 10^12^ vg ml^−1^) with a Nanoject III: AAV1-hSyn-DIO-jGCaMP8f (photometry), AAV1-hSyn-DIO-jGCaMP7s (1p imaging), AAV5-hSyn-DIO-ChR2(H134R)-eYFP, AAV1-hSyn-SIO-stGtACR2-FusionRed, AAV2-eF1a-DIO-taCasp3-TEVp, or AAV2-eF1a-DIO-mCherry (control). Optical fibres (0.2 mm, 0.39 NA), 0.5 mm GRIN lenses (Bruker, 100 μm above injection), microwire arrays (16 ch, 35 μm), and headbars were implanted as required and secured with Metabond and dental cement. Mice recovered ≥3 weeks before behaviour.

### Optogenetics

Bilateral 473 nm light (5 mW per fibre, 20 Hz, 10 ms pulses for ChR2; constant for stGtACR2) was delivered through a rotary joint with TTL control (Feather M4). Free-feeding sessions used PRE/ON/POST 10-min epochs with chow or 60% HFD (Research Diets D12492); intake was measured by pellet weight. Real-time place preference was scored from overhead video in Bonsai. Non-consumption interactions with chow, HFD, or a Lego brick were hand-scored in BORIS.

### Head-fixed licking

Mice (not food/water restricted) licked a capacitive spout dispensing 3 μL of 80% chocolate Boost per lick on a custom 3D-printed wheel rig. Licks were detected with Adafruit FreeTouch on a Feather M4 (25 ms debounce). Bouts required ≥5 licks, ≥0.25 s, and ILI < 1 s. Open-loop blocks delivered 5 s or 95 s stimulation independent of behaviour; closed-loop blocks delivered 1 s of light per lick (re-triggered on each lick).

### Fibre photometry

A tri-colour system (RWD R821) recorded jGCaMP8f at 470 nm with an interleaved 410 nm isosbestic at 20 Hz (20–60 μW). Signals were debleached by biexponential fit to the 410 nm channel, motion-corrected by least-squares regression, downsampled to 5 Hz, and z-scored across the session; peri-event traces used a pre-event baseline. Ghrelin (1 mg kg^−1^), CCK (10 μg kg^−1^), or saline (10 μL g^−1^) were given s.c. in a counterbalanced within-subject design with ≥5 min pre and ≥30 min post recording, followed by FED3 pellet access.

### In vivo electrophysiology

Sixteen-channel tungsten microwire arrays in VP were recorded on Plexon OmniPlex. Spikes were sorted offline (Plexon Offline Sorter; J3 and Davies–Bouldin metrics), yielding 13 single units (analysed) and 28 multi-units. ChR2 responses were tested with 20 × 10 s trains of 465 nm light (20 Hz, 10 ms, ≤5 mW). Units were activated (Wilcoxon signed-rank, ≤5 ms) or inhibited (≤10 ms).

### 1p endoscopic imaging

A Mightex OASIS bundle endoscope (0.5 mm) imaged VP^GABA jGCaMP7s at 20 Hz (470 nm LED, 200–500 μW). Movies were bandpass-filtered, spatially downsampled to 256 × 256, temporally downsampled to 10 Hz, and motion-corrected with TurboReg in CIAtah; sources were extracted with EXTRACT (preprocessing disabled). ΔF/F used the 10th-percentile baseline. Cells were classified as bout-activated, -inhibited, or non-responsive by Wilcoxon signed-rank on bout vs pre-bout ΔF/F with Benjamini–Hochberg FDR < 0.05. For decoding, L2 logistic regression (C = 1.0) was trained on session-z-scored population vectors at each frame from ™3 to +10 s relative to bout onset to discriminate short from long bouts (session-quartile thresholds), using stratified 5-fold CV with balanced accuracy; shuffled bout-label controls were generated, and per-session auROCs were computed over post-onset windows and averaged by mouse.

### Whole-brain c-Fos mapping

Light-sheet imaging of c-Fos in cleared brains from ad libitum vs 12 h–fasted males (n = 8 per group) was performed by Gubra A/S, with counts registered to the GUBRA Multimodal Atlas Framework (Supplementary Table 2).

### Diet-induced obesity and home-cage behavior

Singly housed mice received ad libitum 60% HFD with weekly weights. IPGTT used a 6 h fast and 2 g kg^−1^ glucose i.p., with tail-vein glucose at 0, 15, 30, 60, 90, 120 min (Accu-Chek Guide Me). FED3 devices logged feeding, a 2-armed bandit (reversal every 20 pellets, 3 d), a closed-economy progressive ratio (+1 per pellet, reset after 30 min of no pokes), and an FR1 social operant. TumbleFeeders recorded HFD intake and Pallidus MR1 sensors recorded home-cage activity.

### RNAscope and histology

Multiplex Fluorescent v2 (ACD) on 30 μm fixed-frozen sections used Slc32a1 (319191), Slc17a6 (456751), and DAPI; cells were counted in ImageJ ROIs centred on VP. Perfused brains (4% PFA, 30% sucrose, 30 μm) were stained with chicken anti-GFP (Aves GFP-1020, 1:1,000) where needed and imaged on a Leica DFC7000 widefield. Placements were verified against the Allen CCFv3; off-target animals were excluded.

### Transcriptomic analyses

ABC-Atlas MERFISH (Zhuang-ABCA-1) cells were displayed in CCFv3 with cluster-level mean log2-normalized scRNA-seq expression from the ABC Atlas projected onto each cell by transcriptomic cluster (reference-based, not de novo imputation). snRNA-seq from ARC (GSE276414) and VP (GSE277465) were analysed in Seurat with UMAP, dot plots, and AddModuleScore on curated feeding-receptor gene sets.

### Statistics

Analyses used Python (SciPy, statsmodels, scikit-learn), R (Seurat), and MATLAB (CIAtah, EXTRACT). Paired within-subject comparisons used paired t-tests; between-group used Student’s or Welch’s t-tests, or Mann–Whitney U. Multi-group designs used one-way or mixed RM-ANOVA with Holm–Bonferroni, Bonferroni, or Tukey HSD post-hoc tests; Friedman was used as the non-parametric RM alternative. Normality was assessed with Shapiro–Wilk. Tests were two-tailed, α = 0.05; data are mean ± s.e.m. Exact n, df, test statistics and P values are in figure legends, with effect sizes in Supplementary Table 1.

### AI disclosure

Anthropic Claude (claude-sonnet-4-6) and EdisonScientific assisted with custom Python/MATLAB/R scripts, text drafting, and proofreading. Authors reviewed output and take responsibility for the final article.

## Acknowledgements

This work was supported by the NIH R01 DK136810 (AVK), DK138131 (AVK), DA049924 (MCC), F31 DK138755 (JGW), the Washington University Diabetes Research Center Pilot Award Program (AVK and MCC), the Washington University Nutrition Obesity Research Center Pilot Award Program (AVK), and the Taylor Family Institute for Innovative Psychiatric Research, Washington University School of Medicine. We thank members of the Kravitz and Creed labs for their careful feedback and advice throughout this project.

