## Supplementary material for "Ventral pallidal GABAergic neurons control hedonic feeding and obesity": Figure S1-10

### Supplemental Figure 1:

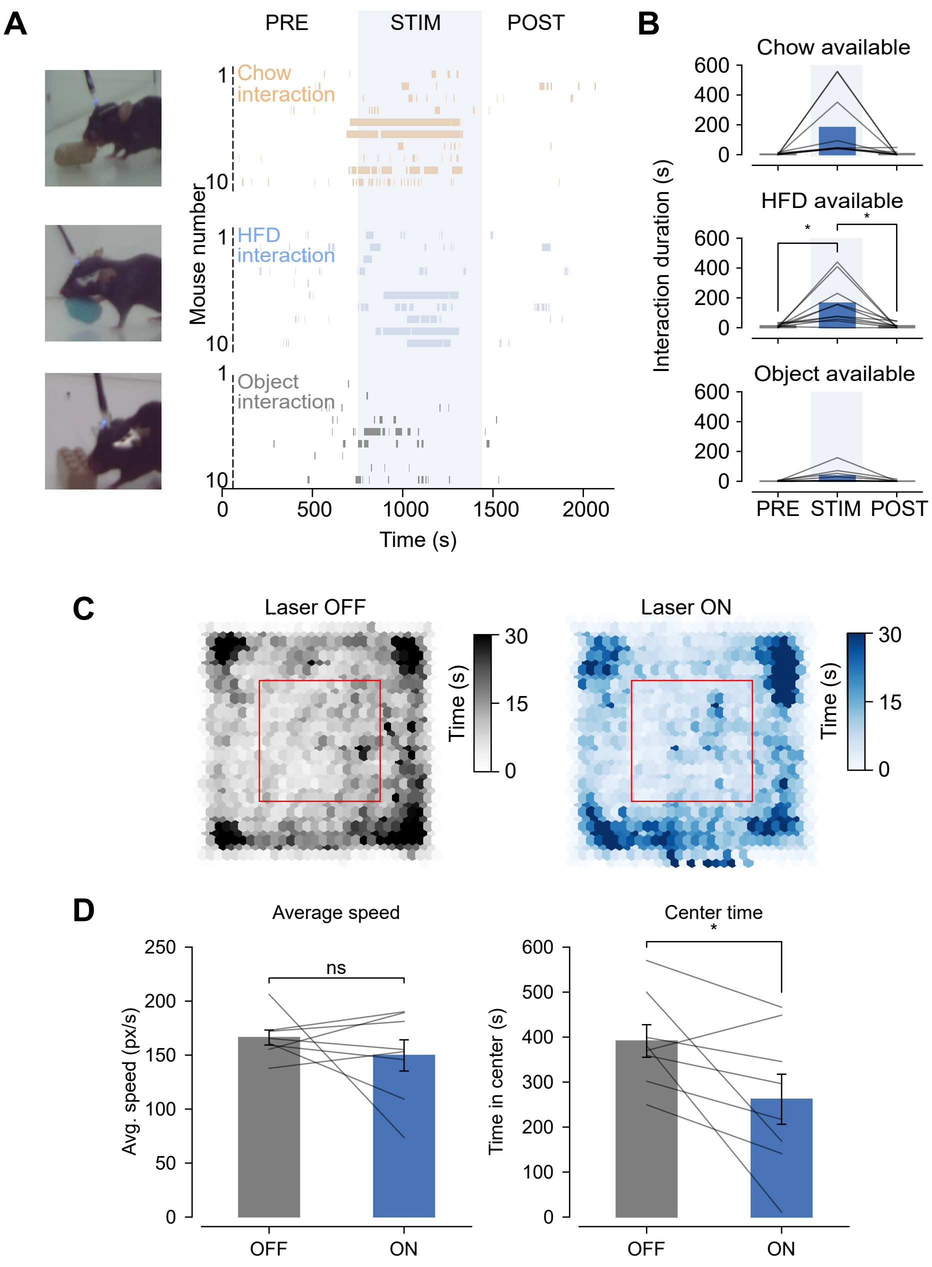


**Supplemental Figure 1. Behavioral ethograms and locomotor tracking during optogenetic VP$^{GABA}$ stimulation.**

**A.** Ethogram raster of interaction of chow, high-fat diet (HFD), and a non-food object (Lego) across pre-laser (PRE), laser-ON (STIM), and post-laser (POST) epochs in VP$^{GABA:ChR2}$ mice.

**B.** Interaction duration during PRE, STIM, and POST epochs for chow, HFD, and non-food object.

**C.** Arena-occupancy heatmaps during all laser OFF and laser ON stimulation sessions. Red box denotes center area for analysis.

**D.** Average locomotor speed and time spent in center during laser OFF and laser ON stimulation sessions. Displayed symbols indicate * = p<0.05, ** = p<0.01, *** = p<0.001, error bar indicates mean ± SEM. For detailed statistics see **Supplemental Table 1**.

**Supplemental Figure 2:**

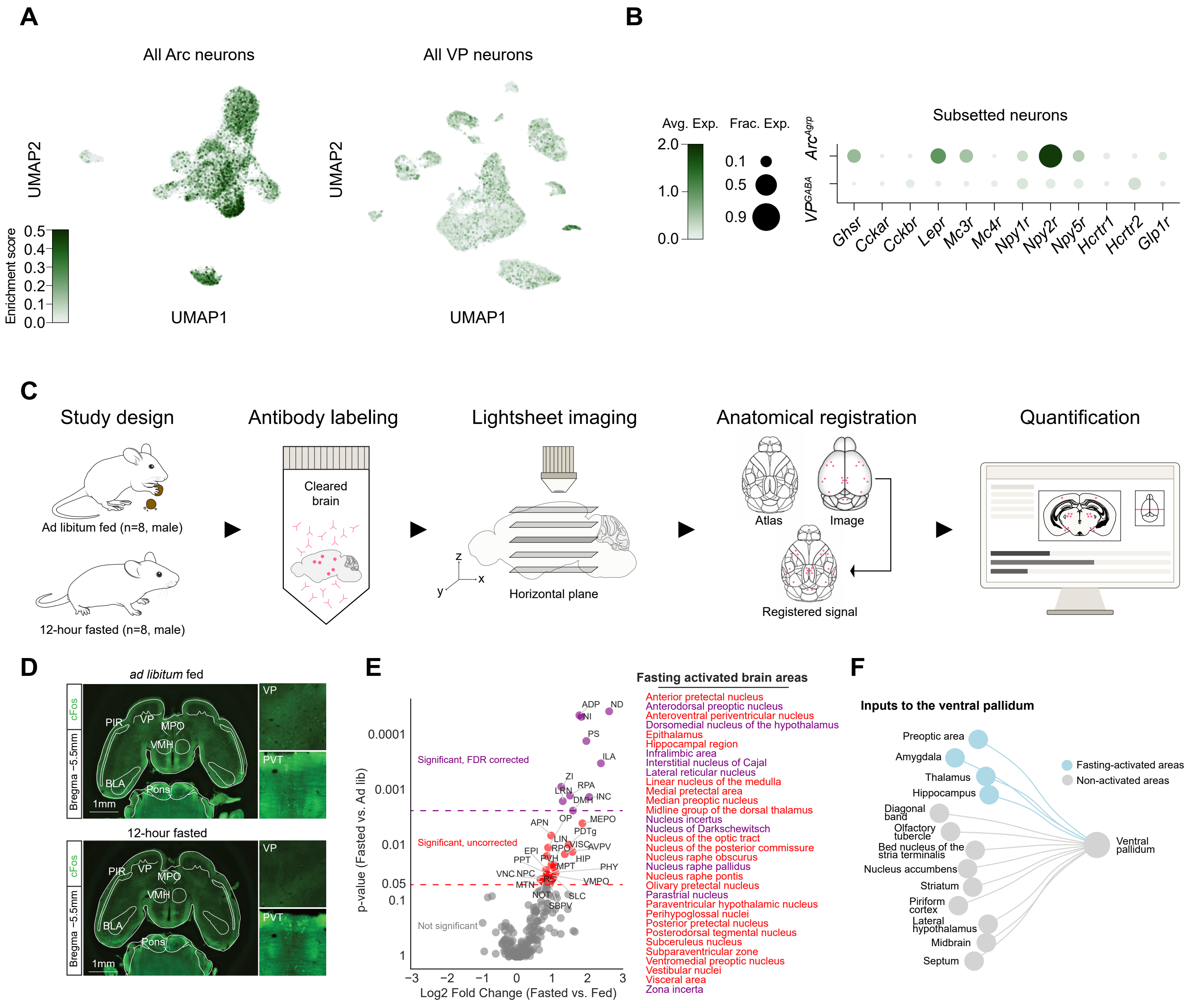


**Supplemental Figure 2. Single cell RNA-sequencing of the VP and whole-brain mapping of fasting-activated inputs to the VP.**
**A.** Dimensionality reduction using Uniform Manifold Approximation and Projection (UMAP) of all arcuate nucleus (Arc) and ventral pallidum (VP) neurons from published single-cell RNA sequencing datasets. Cells are colored by Seurat module score reflecting enrichment of queried feeding genes (*Ghsr*, *Cckar*, *Cckbr*, *LepR*, *Mc3r*, *Mc4r*, *Npy1r*, *Npy2r*, *Npy5r*, *Hcrtr1*, *Hcrtr2*, and *Glp1r*) normalized to a matched, randomly sampled background gene set.
**B.** Dot plot of average and fractional expression of queried feeding genes across all sampled neurons within the $Arc^{AgRP}$ and $VP^{GABA}$ subsetted neurons.
**C.** Experimental design for whole-brain cFos workflow. Ad libitum fed (n=8, male) and 12-hour fasted (n=8, male) mice were processed for whole-brain cFos immunolabeling, tissue clearing, light sheet imaging, anatomical registration, and quantification.
**D.** Representative registered cFos signal in horizontal brain sections, including the ventral pallidum (VP) and paraventricular thalamus (PVT), for ad libitum fed and 12-hour fasted mice.
**E.** Volcano plot of cFos counts by region (log2 fold change, fasted vs. fed) against p-value, with fasting-activated brain areas highlighted (uncorrected and FDR-corrected significance indicated by color).
**F.** Schematic of monosynaptic inputs to the VP, colored by fasting-activated regions.
For detailed statistics regarding cFos expression see **Supplemental Table 2**.

**Supplemental Figure 3:**

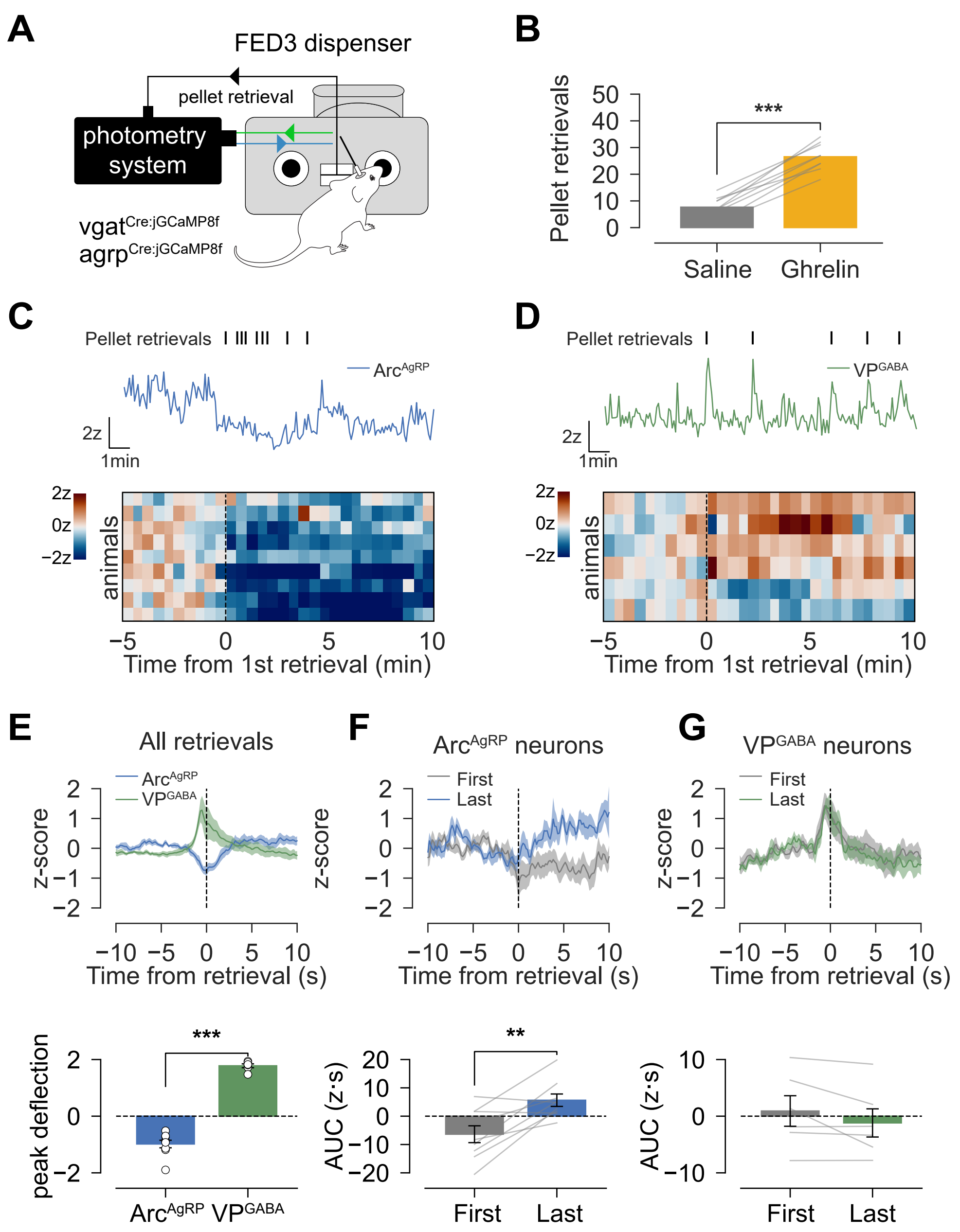


**Supplemental Figure 3. Fiber photometry of Arc$^{AgRP}$ and VP$^{GABA}$ neurons aligned to pellet retrievals dispensed from FED3.**
**A.** Schematic of fiber photometry during FED3 pellet retrieval in VP$^{GABA:jGCaMP8f}$ and Arc$^{AgRP:jGCaMP8f}$ mice.
**B.** Number of pellets taken following Saline or Ghrelin injection.
**C.** Representative Arc$^{AgRP}$ z-scored fluorescence trace with retrieval timestamps; heatmap of all animals displayed below aligned to the first retrieval in session.
**D.** Representative VP$^{GABA}$ z-scored fluorescence trace with retrieval timestamps; heatmap of all animals displayed below aligned to the first retrieval in session.
**E.** Mean z-scored fluorescence aligned to pellet retrieval (-10 to 10s) for Arc$^{AgRP}$ and VP$^{GABA}$ neurons (all-pellet average), averaged by mouse. Peak deflection (highest z-score response) for Arc$^{AgRP}$ and VP$^{GABA}$ neurons plotted below.
**F.** Mean z-scored fluorescence aligned to first and last pellet retrieval within session for Arc$^{AgRP}$ neurons. Average post-event AUC plotted below.
**G.** Mean z-scored fluorescence aligned to first and last pellet retrieval within session for VP$^{GABA}$ neurons. Average post-event AUC plotted below.
Displayed symbols indicate * = p<0.05, ** = p<0.01, *** = p<0.001, error bar indicates mean ± SEM. For detailed statistics see **Supplemental Table 1**.

**Supplemental Figure 4:**

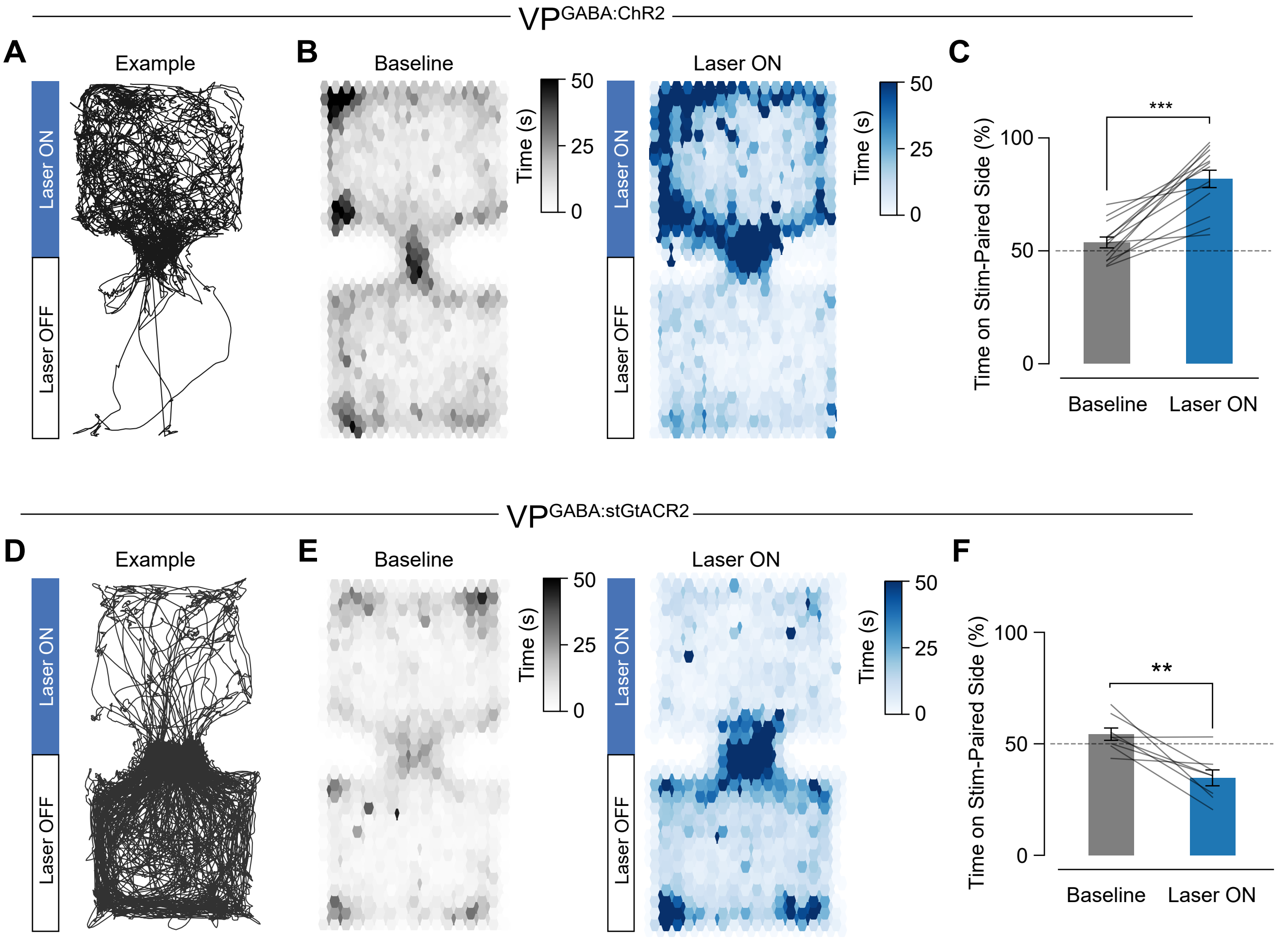


**Supplemental Figure 4. VP$^{GABA}$ optical activation and inhibition drive place preference and avoidance, respectively.**

**A.** Representative movement track of a VP$^{GABA:ChR2}$ mouse during the real-time place preference (RTPP) laser ON session, with the stimulation-paired side indicated.

**B.** Occupancy heatmaps for all baseline (OFF) and laser ON sessions in VP$^{GABA:ChR2}$ mice.

**C.** Percent time spent on the stimulation-paired side, between baseline vs. laser ON sessions in VP$^{GABA:ChR2}$ mice.

**D.** Representative movement track of a VP$^{GABA:stGtACR2}$ mouse during RTPP laser ON session.

**E.** Occupancy heatmaps for all baseline (OFF) and laser ON sessions in VP$^{GABA:stGtACR2}$ mice.

**F.** Percent time spent on the stimulation-paired side, between baseline vs. laser ON sessions in VP$^{GABA:stGtACR2}$ mice.

Displayed symbols indicate * = $p<0.05$, ** = $p<0.01$, *** = $p<0.001$, error bar indicates mean ± SEM. For detailed statistics see **Supplemental Table 1**.

### Supplemental Figure 5:

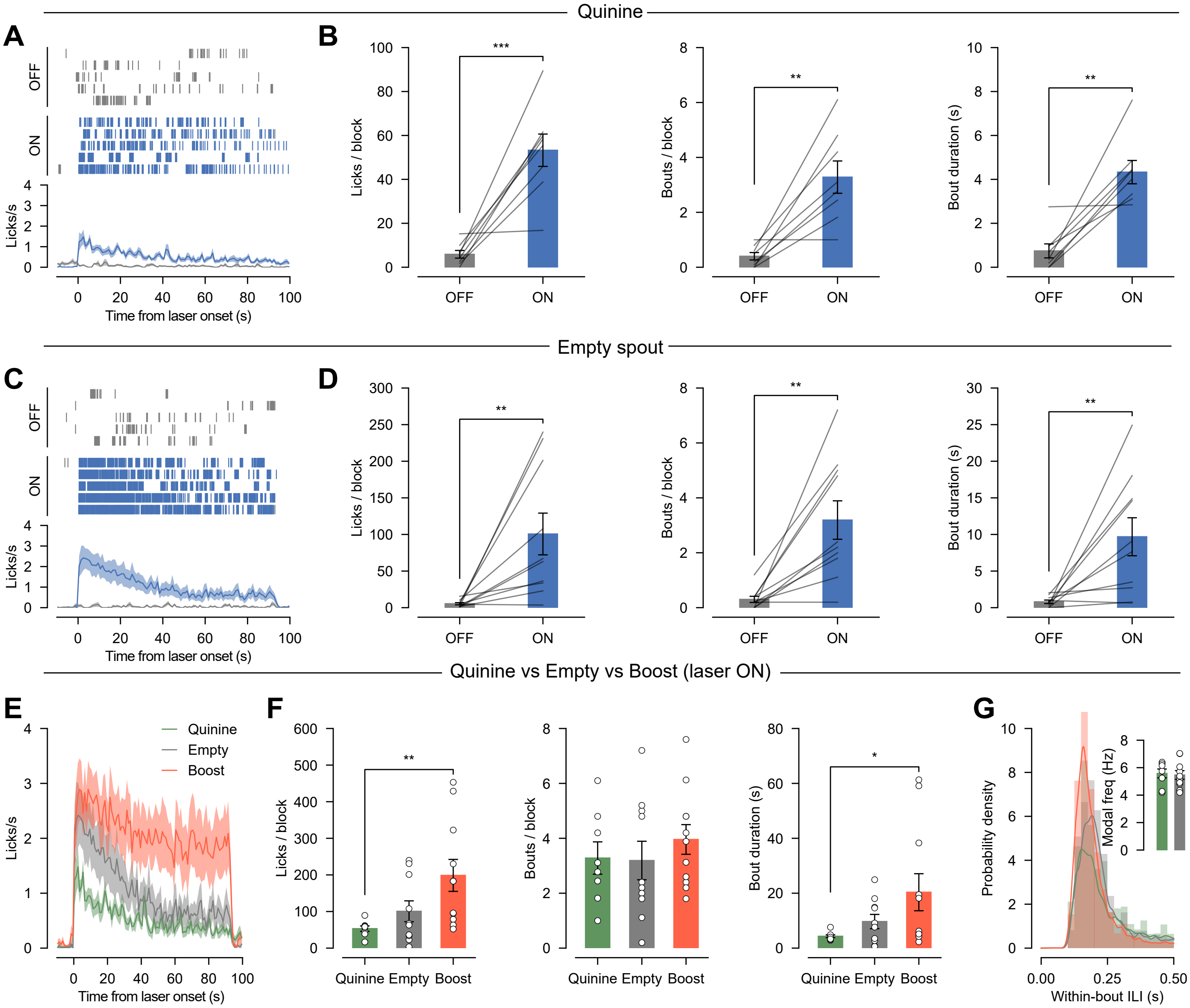


**Supplemental Figure 5. Optogenetic activation of VP$^{GABA}$ neurons drives licking for quinine and an empty spout.**

**A.** Representative per-block lick raster (from a single mouse) and mouse averaged licking rate, plotted by laser OFF vs. ON blocks, for quinine. In laser ON blocks, 20hz stimulation (5mW, 10ms pulse width) was delivered for 95s. Data aligned to laser onset.

**B.** Licking microstructure analysis for quinine, by laser OFF vs. ON blocks. From left to right: licks / block, bouts / block, and average bout duration.

**C.** Representative per-block lick raster (from a single mouse) and mouse averaged licking rate plotted by laser OFF vs. ON blocks, for an empty spout (no reward delivered). In laser ON blocks, 20hz stimulation (5mW, 10ms pulse width) was delivered for 95s. Data aligned to laser onset.

**D.** Licking microstructure analysis for empty spout, by laser OFF vs. ON blocks. From left to right: licks / block, bouts / block, and average bout duration.

**E.** Mouse averaged licking rate during laser ON blocks for quinine, empty spout, and Boost reward. Data aligned to laser onset.

**F.** Licking microstructure analysis during laser ON blocks for quinine, empty spout, and Boost reward. From left to right: licks / block, bouts / block, and average bout duration.

**G.** Within bout distribution of inter-lick interval and mouse averaged modal frequency during laser ON blocks for quinine, empty spout, and Boost reward.

Displayed symbols indicate * = p<0.05, ** = p<0.01, *** = p<0.001, error bar indicates mean ± SEM. For detailed statistics see **Supplemental Table 1**.

**Supplemental Figure 6:**

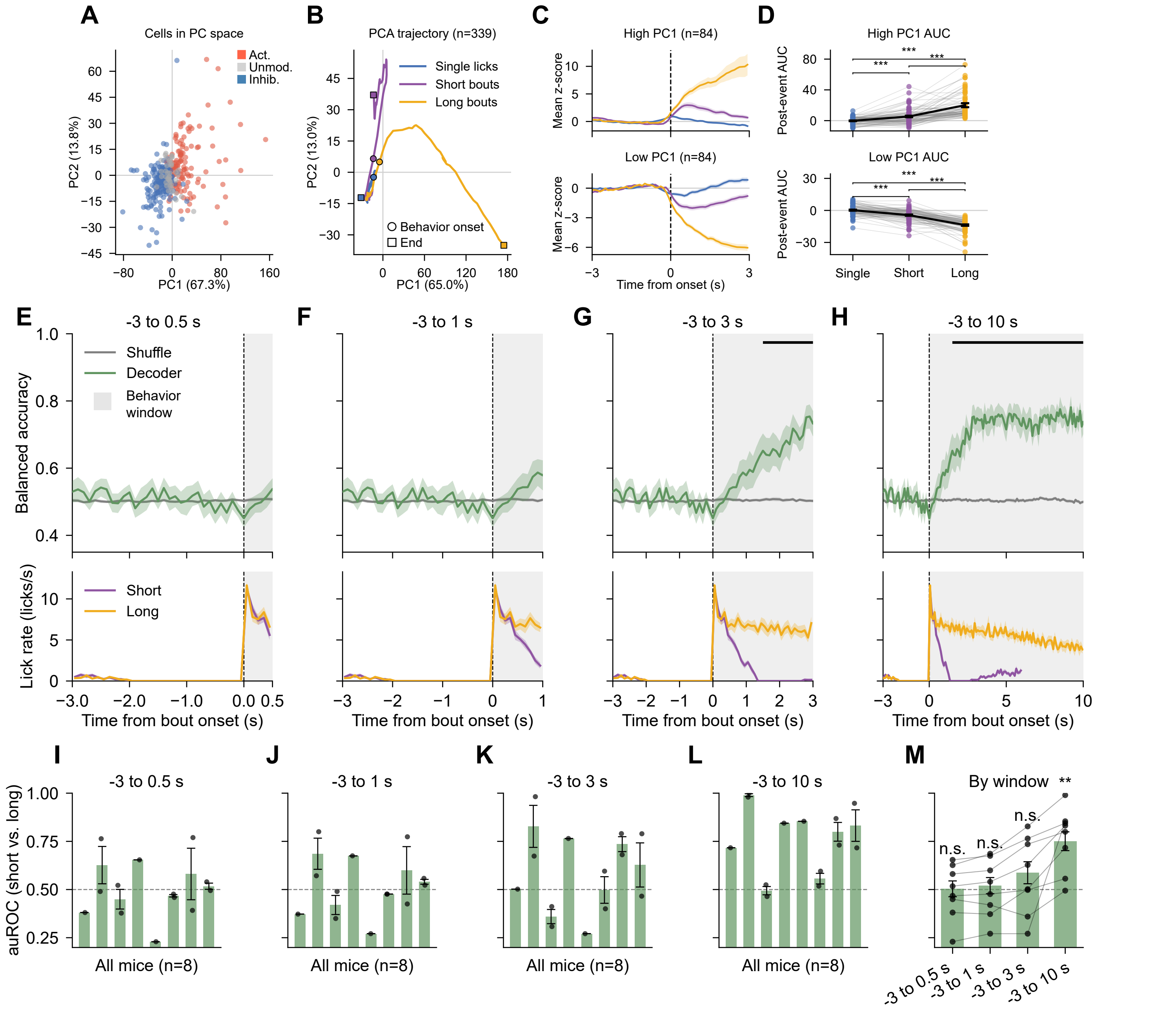


**Supplemental Figure 6. Principal component state dynamics of VP^GABA neurons and linear decoding of VP^GABA population dynamics during self-paced drinking.**

**A.** Z-scored fluorescence traces (-3s to +3s) aligned to lick (or bout) onset of all recorded cells (n=339) plotted along the first two principal components (PC1, 67.3% of variance; PC2, 13.8% of variance). Cells are colored by response classification to licking (activated, unmodulated, inhibited).

**B.** Population trajectories of z-scored fluorescence traces (-3s to +3s) in PC1–PC2 space for single licks, short bouts, and long bouts, from behavior onset to end.

**C.** Cells belonging to the top and bottom 25% coefficients of PC1 eigenvector, plotted by their z-scored fluorescence traces from -3s to +3s from the behavior onset.

**D.** Post-event AUC of top and bottom 25% PC1 eigenvector coefficients during single licks, short bouts, and long bouts.

**E-H.** Balanced accuracy of a linear decoder classifying lick-bout type (short vs. long) from all recorded VP^GABA cells, plotted along label-shuffled data for progressively longer windows from bout onset. Licking rate for short and long bouts shown below.

**I-L.** Per mouse decoder area under Receiver Operator Characteristic (auROC) across all windows. Dashed line indicates chance level auROC.

**M.** Summary of decoder auROC across all windows. Decoder performance significantly exceeded chance only during -3s to +10s window.

Displayed symbols indicate * = $p<0.05$, ** = $p<0.01$, *** = $p<0.001$, error bar indicates mean ± SEM. For detailed statistics see **Supplemental Table 1.**

### Supplemental Figure 7:

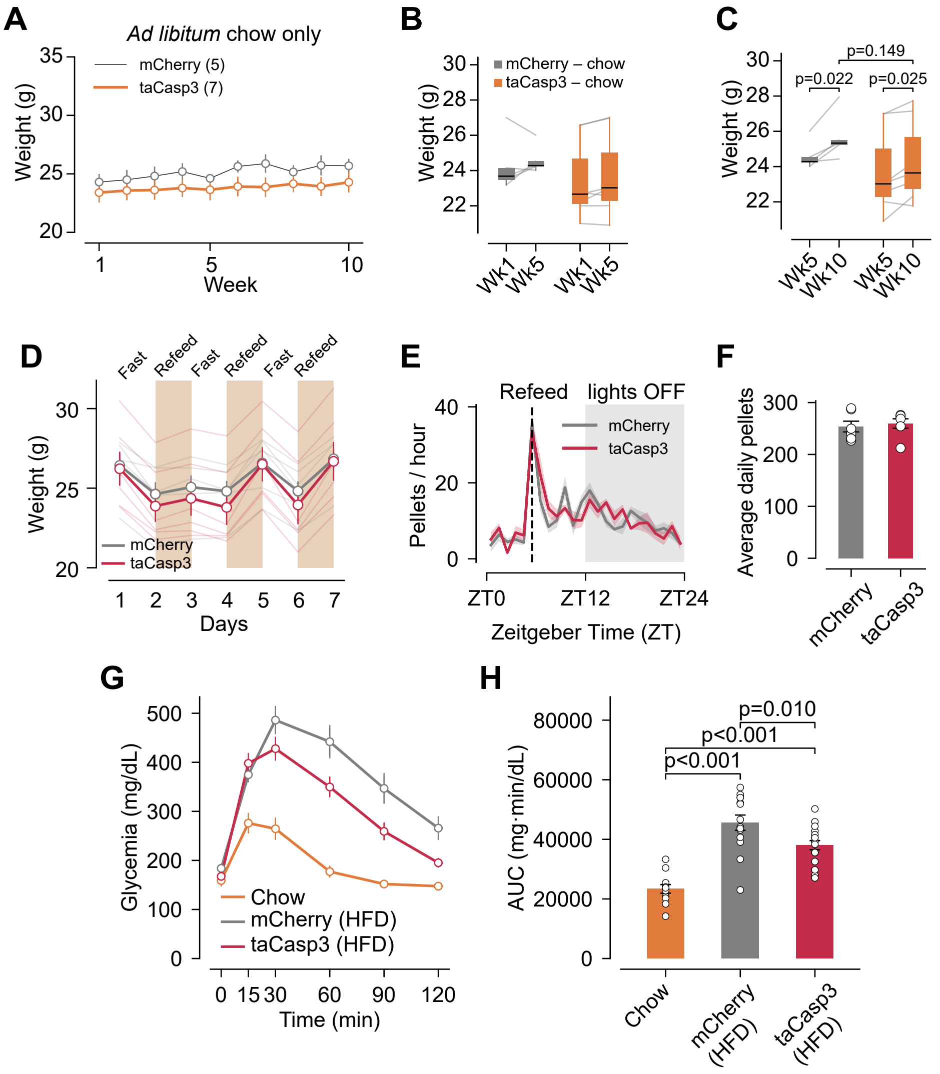


**Supplemental Figure 7. VP$^{GABA}$ ablation leaves homeostatic weight regulation intact, but HFD exposure in VP$^{GABA}$ ablated mice induces hyperglycemia.**
**A.** Body weight trajectories of chow-fed mCherry (n=5) and taCasp3 (n=7, VP$^{GABA}$ ablated) mice across 10 weeks.
**B.** Body weight comparison between week 1 and week 5 of chow-exposed animals.
**C.** Body weight comparison between week 5 and week 10 of chow-exposed animals.
**D.** Body weight across 7 days of alternating fast and refeed (FED3 access) cycles for chow-fed mCherry vs. taCasp3 expressing animals.
**E.** Averaged pellet-retrieval rate per hour across the circadian cycle aligned to each refeeding period.
**F.** Average daily pellets for mCherry and taCasp3 animals, during refeeding periods.
**G.** Glycemia measurements during glucose tolerance test (GTT) for chow only, mCherry HFD exposed, and taCasp3 HFD exposed mice. Glucose injection delivered at t=0 min.
**H.** GTT AUC from 0min to 120min.
Displayed symbols indicate * = p<0.05, ** = p<0.01, *** = p<0.001, error bar indicates mean ± SEM. For detailed statistics see **Supplemental Table 1**.

### Supplemental Figure 8:

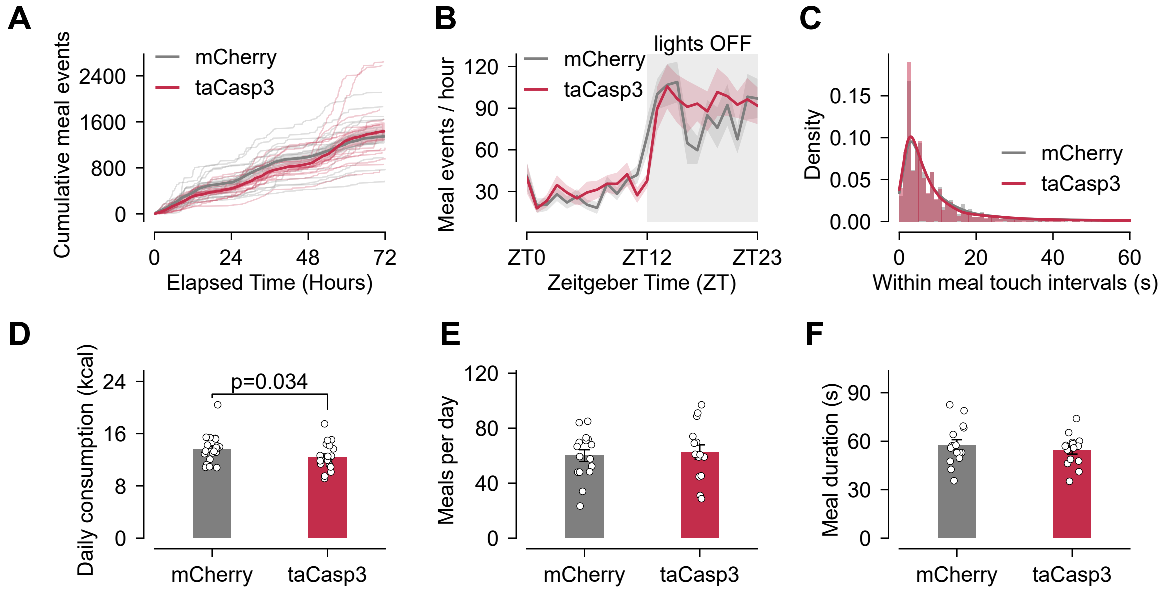


**Supplemental Figure 8. VP$^{GABA}$ ablation does not disrupt circadian feeding patterns for HFD.**

**A.** Self-paced HFD feeding of singly housed mCherry vs taCasp3 expressing mice measured by cumulative hopper touches (capacitive sensing, TumbleFeeder device) over 72 hours in the home cage. Mice were exposed to HFD for 5 weeks prior to this measurement.

**B.** Daily average of hopper touches per hour plotted across circadian cycle.

**C.** Hopper touches were classified as “meals” with cut-off criteria. Probability distribution of within meal touch intervals of all meals during 72-hour period.

**D-F.** Daily HFD consumption, meals per day, and meal duration for mCherry vs taCasp3 expressing mice. Error bar indicates mean ± SEM. For detailed statistics see **Supplemental Table 1**.

### Supplemental Figure 9:

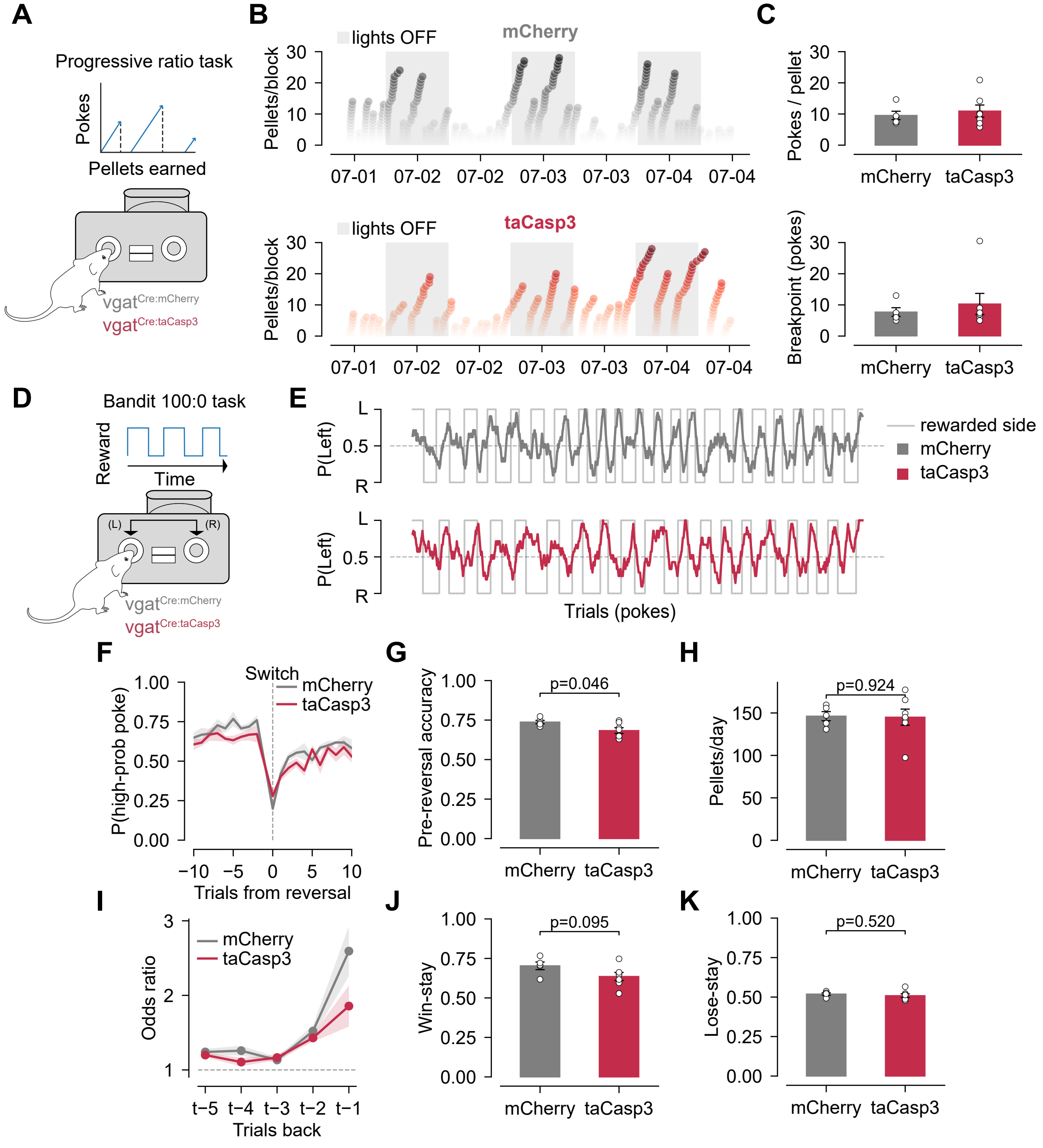


**Supplemental Figure 9. Effect of VP$^{GABA}$ ablation on operant reward seeking and reversal learning.**
**A.** Schematic of the progressive-ratio operant task. HFD-naïve mice were tested.
**B.** Representative pellets-earned rasters across days.
**C.** Pokes per pellet and breakpoint (pokes before block reset) in mCherry vs taCasp3 expressing mice.
**D.** Schematic of the 100:0 probabilistic-reversal ('bandit') task. HFD-naïve mice were used.
**E.** Representative P(left) choice behavior with reward-side contingency.
**F.** Probability of a high-probability poke aligned -10 and +10 trials from reversal.
**G.** Average pre-reversal accuracy for mCherry vs taCasp3 expressing mice.
**H.** Average daily pellets for mCherry vs taCasp3 expressing mice.
**I.** Logistic-regression odds ratios for reward at trial lags t-5 to t-1.
**J.** Average win-stay rate for mCherry vs taCasp3 expressing mice.
**K.** Average lose-stay rate for mCherry vs taCasp3 expressing mice.
Error bar indicates mean ± SEM. For detailed statistics see **Supplemental Table 1**.

**Supplemental Figure 10:**

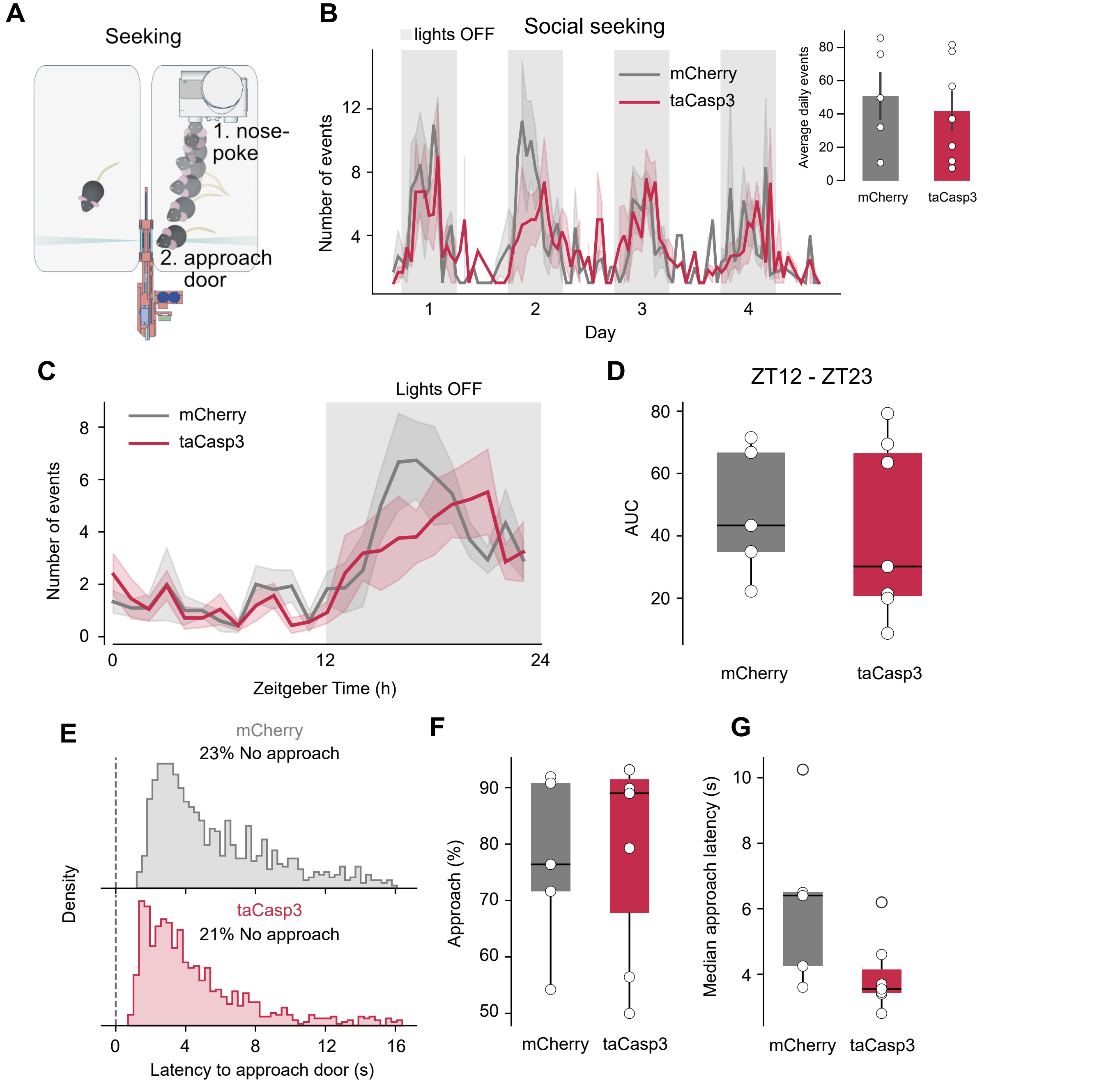


**Supplemental Figure 10. Social seeking behaviors remain intact following VP$^{GABA}$ ablation.**
**A.** Schematic of operant social seeking task. HFD-naïve mice were used.
**B.** Number of social-seeking events across days. Inset displays average daily events for mCherry vs taCasp3 expressing mice.
**C.** Circadian plot of social-seeking events per hour, averaged across 4 days.
**D.** AUC of social-seeking events during the dark phase (ZT12–ZT23).
**E.** Distribution of latency to approach the social-access door, with the percentage of trials with no approach indicated.
**F.** Percentage of trials with approach to door for mCherry vs taCasp3 expressing mice.
**G.** Median approach to door latency for mCherry vs taCasp3 expressing mice.
Error bar indicates mean ± SEM. For detailed statistics see **Supplemental Table 1**.
