## Supplemental Methods for "Ventral pallidal GABAergic neurons control hedonic feeding and obesity"

**Subjects.** Adult male and female mice (≥8 weeks old at surgery; ≥3–4 weeks of viral expression before behavior, ~12 weeks at testing) were used. Both male and female mice were used but cohorts were not sex-balanced and were underpowered to treat sex as a categorical factor; male and female data were pooled. Group sizes are reported in the main text and figure legends. *Vgat*-IRES-Cre (Jax #028862) and *Agrp*-IRES-Cre (Jax # 012899) mice were obtained from The Jackson Laboratory and bred with C57BL/6J wildtype mice to produce experimental offspring. All procedures were approved by the IACUC of Washington University in St. Louis, School of Medicine.

**Viral vectors.** Calcium indicators: AAV1-hSyn-DIO-jGCaMP8f (Addgene #162379, fiber photometry) and AAV1-hSyn-DIO-jGCaMP7s (Addgene #104491, 1-photon endoscopic imaging). Excitatory opsin: AAV5-hSyn-DIO-ChR2(H134R)-eYFP (UNC Viral Vector Core). Inhibitory opsin: AAV1-hSyn-SIO-stGtACR2-FusionRed (Addgene #105677). Chronic cell-type-specific ablation: AAV2-eF1a-DIO-taCasp3-TEVp (UNC Viral Vector Core), with AAV2-eF1a-DIO-mCherry (UNC) as the serotype-matched fluorophore control.

**Intracranial stereotaxic surgery.** Mice were anesthetized with isoflurane (5% induction, 1–2% maintenance). VP coordinates from bregma were AP +0.1, ML ±1.5, DV −4.95 mm. Viral vectors (diluted to 6–8 × $10^{12}$ vg/mL) were delivered through glass capillaries with a Drummond Nanoject III at 10 nL/s in ≥5 cycles to a final volume of 250–300 nL. Injections were unilateral for photometry and endoscopic imaging and bilateral for optogenetics and ablation. For head-fixation or optical stimulation, an optical fiber cannula (RWD #907-03007-00; 6 mm, 0.2 mm core, 0.39 NA) and a custom stainless-steel headbar (2 mm, Ponoko) were implanted at the time of viral injection. For endoscopic imaging, a 0.5 mm GRIN lens (Bruker #1050-004600) was implanted 100 µm above the injection site. All implants were secured with C&B-Metabond (Parkell) followed by black Lang Ortho-Jet dental cement (Lang Dental). Mice recovered for ≥3 weeks before behavior.

**Free-moving optogenetic experiments.** For ChR2 activation experiments, blue light (473 nm DPSS laser) was delivered bilaterally through a 50:50 split rotary joint (Doric FRJ_1x2i_FC-2FC) and patch cords (RWD #807-00061-00), with TTL timing (20 Hz, 10 ms pulses) controlled by a Feather M4 microcontroller (Adafruit). Laser power was verified to be 5mW at each fiber tip before each session. **Free-moving food intake measurements** consisted of three 10-min epochs (PRE, ON, POST), with light pulsed continuously during the ON period, and not during the other periods. A single pellet of chow (PicoLab® Rodent Diet 20, LabDiet) or 60% HFD (Research Diets #D12492) was available throughout all three periods, and the weight of the pellet was weighed at the end of each epoch to determine consumption. **Real-time place preference (RTPP)** experiments were conducted in a 12” x 24” arena split into two equal zones (**Fig S4**), with a short hallway between them. Position was tracked online by a top-mounted camera, and Bonsai[1] was used to trigger the laser on when the mouse was on a prespecified side. Preference was quantified as the fraction of session time in each zone. Head fixed experiments used the parameters above**,** for ChR2 activation.

**Manual annotation of behavior videos during optogenetic stimulation.** To measure non-consumption interactions with standard chow, HFD, or an inedible object (plastic Lego brick), we used Behavioral Observation Research Interactive Software (BORIS)[2] to manually score events from video recordings. Non-consumption interactions were defined as events in which the animal retrieved the food item or object in its mouth and carried or manipulated it without ingestion. Interaction bouts were aligned to the same three 10-minute epoch structure used for freely moving optogenetics consumption measurements (PRE, STIM, POST), where stimulation was delivered exclusively during the ON epoch.

**Fiber photometry recordings.** Fiber photometry recordings were acquired using a tri-color multichannel system (RWD R821). jGCaMP8f was excited at 470 nm; motion artifacts and fluorophore bleaching were corrected using an isosbestic 410 nm channel that was acquired during interleaved frames. Excitation power at the fiber tip was 20–60 µW. Signals were sampled at 20 Hz and synchronized with behavior via TTL pulses. Raw photometry signals were processed using custom Python scripts. Prior to analysis, recordings were manually inspected using an interactive GUI (custom Python code) to identify and retain only artifact-free epochs (e.g., excluding periods of severe movement-induced fiber disturbance, especially around injections). Photobleaching correction was performed by fitting a biexponential decay to the 410 nm isosbestic reference channel, which was then used to generate a fitted baseline for the 470 nm calcium signal via a least-squares regression. The motion corrected and debleached fluorescence signal was downsampled to 5 Hz and z-scored by subtracting the mean and dividing by the standard deviation computed across the entire recording session. To construct peri-stimulus photometry traces (for injections or pellet retrievals), fluorescence values were z-scored relative to a pre-stimulus baseline period. For comparison across animals and conditions, area under the curve (AUC) was computed over defined peri-event windows around the injection timing. For pharmacology experiments, ghrelin (1 mg/kg), CCK (10 µg/kg), or matched-volume 0.9% saline (10 µL/g) were administered subcutaneously in the home cage while animals were tethered to the photometry system. Drug and vehicle injections were counterbalanced within-subject design and were performed during the light cycle. Photometry was recorded for ≥5 min before and ≥30 min after injection. After the 30 min post-injection period, animals were given access to a pellet dispenser (Feeding Experimental Device 3, FED3[3]) to record calcium dynamics around pellet consumption and to confirm the expected effects of ghrelin on intake.

**In vivo electrophysiology.** Chronic extracellular recordings used 16-channel tungsten microwire arrays (35 µm diameter; Innovative Neurophysiology) implanted in VP at the time of ChR2 injection (coordinates as above). After ≥2 weeks of recovery and expression, signals were amplified, filtered, and recorded on a Plexon OmniPlex system. Spikes were sorted offline in Plexon Offline Sorter; isolation was judged by waveform consistency, ISI distributions, and J3 and Davies–Bouldin metrics, yielding 13 well-isolated single units and 28 multi-units. Only the 13 single units were included for further analysis. Blue-light stimulation (Plexon PlexBright 465 nm LED, up to 5 mW, 20 Hz, 10 ms pulses; twenty 10-s trains) was used to characterize $VP^{GABA}$ responses. Peri-event-stimulus-histograms (PSTHs) were built in NeuroExplorer v5 (Plexon) aligned to LED pulse onset, with 1 ms bins. Units were classified as activated if spiking increased significantly within 5 ms of pulse onset (Wilcoxon signed-rank), and as inhibited if spiking decreased significantly within 10 ms of pulse onset.

**Whole-brain light sheet microscopy of cFos positive nuclei.** Whole-brain mapping of cFos-positive nuclei was performed by Gubra A/S (Copenhagen, Denmark) using light-sheet fluorescence microscopy. Male mice were given *ad libitum* access to food or fasted overnight (12 h) prior to perfusion (n=8 per group). Tissue clearing, immunolabeling, imaging, and cell detection were performed using Gubra's pipeline[4]. Brain-wide cFos counts were registered to the GUBRA 3D Mouse Brain Atlas Framework, and regions showing differential expression between sated and fasting conditions are reported in Supplemental Table 2.

**Head-fixed optogenetic experiments**. A custom head-fixation apparatus was constructed using 3D-printed components, and a capacitive lickometer, modeled after[5], design files here: https://github.com/KravitzLabDevices/Head_fixed_wheel). Mice were not food or water restricted and all sessions were conducted in the light cycle. Prior to experimental sessions, animals underwent habituation and training to perform self-paced licking until consumption stabilized (typically 1 week). In open-loop stimulation paradigms, each lick triggered a 3µL delivery of 80%-diluted chocolate Boost via opening a solenoid. Individual lick timestamps were detected via capacitive touch sensing (Adafruit Freetouch Library) and digitized by an Arduino Feather M4 (Adafruit Industries). A 25ms debounce was applied in software to exclude spurious double-detections. Lick bouts were defined using a sliding 1-s inter-lick interval (ILI) threshold: consecutive licks separated by less than 1 s were grouped into the same bout, while licks separated by more than 1 s were assigned to a new bout. Bouts were further required to meet minimum criteria of ≥5 licks and ≥0.25 s duration. Microstructure metrics including bout duration, bouts per session, licks per bout, and modal within-bout lick frequency (estimated as the reciprocal of the mode of the inter-lick interval distribution via kernel density estimation) were computed for each session. A wheel encoder also recorded movements on the wheel, which were converted to instantaneous velocity offline. Behavioral acquisition and hardware synchronization were performed using Bonsai[1] with an Arduino UNO running the Firmata protocol. Optogenetic stimulation or inhibition experiments paired the licking with laser stimulation as follows: Open-loop protocol: 20 Hz stimulation (for ChR2-mediated activation) or constant stimulation (for stGtACR2-mediated inhibition) in fixed 5 s or 95 s blocks independent of behavior (Fig. 2B). Closed-loop protocol: each lick triggered 1 s of stimulation, with the 1 s window resetting on every subsequent lick (i.e., light stayed on until ≥1 s elapsed without a lick). Stimulation blocks were interleaved with no-stimulation blocks.

**1-photon endoscopic calcium imaging.** Head-fixed imaging used a Mightex OASIS Implant fiber bundle endoscopic imaging system (Mightex #FBR-060-30K-100-03, 0.5mm diameter). Cre-dependent jGCaMP7s was expressed in $VP^{GABA}$ neurons and was excited using a 470 nm LED (Mightex BLS-Series, constant illumination, 200-500 µW) with images acquired at 20 Hz with a PCO Edge camera (Excelitas) and synchronized to behavioral timestamps in PolyScan4 acquisition software. Raw .TIF stacks were preprocessed in CIAtah[6] with spatial bandpass filtering, spatial downsampling to 256 × 256 px, temporal downsampling to 10 Hz, and TurboReg motion correction[7]. Sources were extracted from the preprocessed movies with EXTRACT[8]) for GPU-accelerated (NVIDIA RTX5090) source extraction and signal demixing. EXTRACT was run with preprocessing disabled (preprocessing was handled entirely by CIAtah upstream) to avoid redundant spatial filtering. Extracted fluorescence traces were normalized to ΔF/F using a baseline computed as the 10th percentile fluorescence across the session. Individual neurons were classified as consumption-activated, consumption-inhibited, or non-responsive based on Wilcoxon signed-rank tests comparing mean ΔF/F during licking bouts to the pre-bout baseline ($p < 0.05$ after Benjamini-Hochberg false discovery rate correction), and by the sign of the mean response[9]. Behavioral events were synchronized with imaging using the Mightex PolyScan4 acquisition software and Mightex PolyEcho Intelligent Control Module.

A linear classifier was trained to distinguish short from long bouts (as defined by the global session quartile thresholds) using EXTRACTed cells obtained from 1-photon calcium imaging. Calcium traces were normalized using session-wide z-scoring prior to decoding, such that absolute activity differences between bout types were preserved rather than removed by per-trial normalization. At each time point from -3 to +10s relative to the bout onset, we constructed the population activity vector – the z-scored fluorescence of all simultaneously recorded neurons at the imaging frame closest to that time – and used it as the feature vector for classification. A logistic regression classifier (L2 regularization, C=1.0) was trained to discriminate short from long bouts independently at each time point using stratified 5-fold cross-validation, which yielded a balanced accuracy score for each time point, to account for class imbalance between short and long bout trials. Accuracy curves were averaged across sessions (mean ± SEM), with each session's curve resampled to a common time grid before averaging. A shuffle control was generated by permuting bout-type labels (short or long). To obtain a summary statistic per session, population activity was averaged across a range of post bout onset windows (up to 10s) per trial and classified using the same 5-fold procedure, to obtain per-session area under Receiver Operator Characteristic curves (auROC), averaged by mouse.

**Diet-induced obesity model.** Individually housed mice were given ad libitum HFD (60% of calories from fat, Research Diets #D12492). Animals were singly housed and weights were recorded weekly. Glucose tolerance testing was performed by an intraperitoneal glucose tolerance test conducted after a 6-hour fast. Fasting blood glucose was measured via tail vein blood sampling using a glucometer (Accu-Chek Guide Me; upper detection limit 600 mg/dL), followed by an intraperitoneal injection of glucose (2g/kg, 20% solution in sterile saline). Blood glucose was then measured at 15, 30, 60, 90, and 120 minutes post-injection to assess glucose clearance.

**Home-cage behavioral tasks.** Feeding was monitored with FED3 devices[3], which dispense single pellets based on nose-pokes, logging poke and retrieval timestamps. SD-card data were analyzed in custom Python to compute cumulative intake, meal patterns, and bout structure across the light and dark cycles. FED3 was also programmed to administer a 2-armed bandit task, in which an active poke resulted in a pellet and the other resulted in a 10-second timeout. This task was run for 3 days, where the active poke switched between left and right every 20 pellets. FED3 was also used to administer a closed-economy progressive ratio task. Here, each earned pellet increased the poke requirement for the next pellet by 1. If the mouse withheld poking for 30 minutes, the poke requirement reset to 1. Finally, FED3 was used to administer an FR1-based social operant task, in which each poke on the FED3 would open a gate allowing interactions with a conspecific in a neighboring cage as described in[10]. "TumbleFeeders" were used to measure circadian intake patterns of HFD intake. The TumbleFeeders were filled with HFD, and logged touches on the front of the hopper through capacitive touch sensing. Pallidus MR1 devices (MCCI, Ithaca, NY) were used to sense home-cage activity for circadian activity analysis.

**In situ hybridization (RNAscope).** Fluorescent in situ hybridization was performed on 30 µm fixed frozen brain tissue using the RNAscope Multiplex Fluorescent v2 assay (Advanced Cell Diagnostics) as per manufacturer's instructions. Probes targeting *Slc32a1* (Vgat; ACD 319191), *Slc17a6* (Vglut2; ACD 456751), and a DAPI counterstain were used to characterize the cellular composition of the VP injection site. Sections were imaged on a confocal microscope (Leica Stellaris SP5) at 20× magnification as single-tile acquisitions. Transcript-positive cells and co-expressing cells were manually counted in ImageJ within a defined region of interest centered on the VP.

**Histology and immunofluorescence.** Mice were transcardially perfused with PBS followed by 4% PFA. Brains were post-fixed overnight in 4% PFA and cryoprotected in 30% sucrose/PBS until they sank. Coronal sections (30 µm) were cut on a cryostat and collected in PBS. ChR2-eYFP signal was amplified with chicken anti-GFP (Aves Labs #GFP-1020, 1:1000); other fluorophores were imaged directly. Sections were mounted and coverslipped with Fluoromount DAPI and imaged on a Leica DFC7000-equipped widefield microscope.

Viral expression and fiber/GRIN lens placement were verified against the Allen Mouse Brain Reference Atlas; mice with missed injections or off-target placements were excluded.

**Reference-based projection of scRNA-seq cluster signatures onto MERFISH cells.** Spatial transcriptomic data from the ABC-Atlas MERFISH brain Zhuang-ABCA-1 were used for anatomical visualization[11]. Cells were displayed in Allen CCFv3 space using registered three-dimensional coordinates and transcriptomic cluster annotations. Cells within and around the VP and arcuate hypothalamic nucleus were visualized as 150-µm maximum projections, with anatomical boundaries overlaid for reference. Cluster-level gene expression signatures were generated from the ABC Atlas single-cell RNA-seq dataset[12]. Log2-normalized expression matrices were combined with cell-level taxonomy metadata, and cells were grouped by transcriptomic cluster, donor and sequencing chemistry. For each cluster–donor–chemistry group, mean log2-normalized expression and detection rate, defined as the fraction of cells with non-zero expression, were calculated for each gene. Groups passing the minimum cell threshold were averaged across sequencing chemistries and donors to obtain a final mean-expression profile and detection-rate profile for each transcriptomic cluster. For spatial projection, each MERFISH cell was assigned the mean scRNA-seq expression value of the selected gene corresponding to its transcriptomic cluster. The resulting maps therefore represent reference-based cluster-level expected expression projected onto spatially registered MERFISH cells, rather than direct per-cell MERFISH measurements or de novo imputation from the MERFISH expression matrix. For display, expression values were scaled independently for each gene.

**scRNA-seq dataset analysis** from public scRNA-seq. Publicly available single-nucleus RNA sequencing datasets from the ARC (GSE276414)[13] and VP (GSE277465)[14] were accessed from GEO. UMAP was used for dimensionality reduction and cluster visualization, and dot plots showed gene expression across annotated clusters. Module scores for curated feeding-related receptor gene sets were computed with Seurat's AddModuleScore across Arc and VP cell populations. Dot plots displayed expression of genes of interest across annotated cell clusters. Module scores were computed for curated gene sets using the Seurat AddModuleScore function across ARC and VP cell populations.

**Statistical analysis.** Analyses were performed in Python (SciPy, statsmodels, and scikit-learn), with single-cell RNA-sequencing analysis in R (Seurat) and calcium-imaging source extraction in MATLAB (CIAtah, EXTRACT). Paired, within-subject comparisons (e.g., laser OFF vs ON, stimulation vs baseline, and week-to-week body weight) used paired t-tests; unpaired, between-group comparisons (e.g., mCherry vs taCasp3) used independent-samples (Student's) t-tests, Welch's t-test when variances were unequal, or Mann–Whitney U tests where indicated. Multi-group comparisons used one-way or mixed-design repeated-measures ANOVA, or one-way ANOVA for independent groups, followed by pairwise post-hoc tests with Holm–Bonferroni, Bonferroni, or Tukey HSD correction; normality was assessed with the Shapiro–Wilk test, and the Friedman test was used as the non-parametric repeated-measures alternative where appropriate. Summary data are mean ± SEM unless stated otherwise; tests were two-tailed unless noted, with $\alpha = 0.05$. Test statistics, exact sample sizes, degrees of freedom, and p-values are reported in the figure legends, and full statistical detail, including effect sizes, is provided in **Supplemental Table 1**. Experimenters were not blinded to conditions.

**Declaration of generative AI and AI-assisted technologies in the writing process.** Anthropic Claude (claude-sonnet-4-6) was used to help generate custom Python, MATLAB, and R analysis scripts and to assist in drafting the text. The authors reviewed and refined all AI-generated content and take full responsibility for the accuracy of the published article.
