## Supplemental Table 1 for "Ventral pallidal GABAergic neurons control hedonic feeding and obesity"

| Figure | Measure | Test | Comparison | n (M/F) | df | Test statistic | p-value | Post-hoc p-values | Effect size |
|---|---|---|---|---|---|---|---|---|---|
| **Figure 1. Arc$^{AgRP}$ and VP$^{GABA}$ neurons activate distinct feeding drives.** | | | | | | | | | |
| 1B | Arc$^{AgRP}$ chow consumption | One-way RM-ANOVA + Bonferroni post-hoc | PRE vs STIM vs POST | 6 (3M/3F) | F(2,10) | F = 65.20 | 1.8e-06 | PRE-STIM 6.1e-4<br>STIM-POST 8.8e-4<br>PRE-POST 0.102 | η$^2$p = 0.929; STIM-vs-PRE d_z = -3.94 |
| 1C | Arc$^{AgRP}$ HFD consumption | One-way RM-ANOVA + Bonferroni post-hoc | PRE vs STIM vs POST | 6 (3M/3F) | F(2,10) | F = 35.69 | 2.8e-05 | PRE-STIM 3.6e-3<br>STIM-POST 0.013<br>PRE-POST 3.9e-3 | η$^2$p = 0.877; STIM-vs-PRE d_z = -2.70 |
| 1D | Arc$^{AgRP}$ chow prestimulation | Paired t-test | laser OFF vs ON (STIM) (pre-access) | 6 (3M/3F) | t(5) | t = -6.442 | 1.3e-03 | — | Cohen's d_z = -2.63 |
| 1D | Arc$^{AgRP}$ HFD prestimulation | Paired t-test | laser OFF vs ON (STIM) (pre-access) | 6 (3M/3F) | t(5) | t = -5.317 | 3.2e-03 | — | Cohen's d_z = -2.17 |
| 1F | VP$^{GABA}$ chow consumption | One-way RM-ANOVA + Bonferroni post-hoc | PRE vs STIM vs POST | 12 (5M/7F) | F(2,22) | F = 5.595 | 0.011 | PRE-STIM 0.093<br>STIM-POST 0.135<br>PRE-POST 0.288 | η$^2$p = 0.337; STIM-vs-PRE d_z = -0.72 |
| 1F | VP$^{GABA}$ chow consumption — Males [sex-split] | One-way RM-ANOVA + Bonferroni post-hoc | PRE vs STIM vs POST | 5 (5M/0F) | F(2,8) | F = 0.961 | 0.423 (ns) | ON-vs-PRE 0.981 | η$^2$p = 0.194; STIM-vs-PRE d_z = -0.50 |
| 1F | VP$^{GABA}$ chow consumption — Females [sex-split] | One-way RM-ANOVA + Bonferroni post-hoc | PRE vs STIM vs POST | 7 (0M/7F) | F(2,12) | F = 4.805 | 0.029 | ON-vs-PRE 0.213 | η$^2$p = 0.445; STIM-vs-PRE d_z = -0.83 |
| 1G | VP$^{GABA}$ HFD consumption | One-way RM-ANOVA + Bonferroni post-hoc | PRE vs STIM vs POST | 12 (5M/7F) | F(2,22) | F = 31.23 | 3.8e-07 | PRE-STIM 5.1e-4<br>STIM-POST 3.9e-4<br>PRE-POST 0.358 | η$^2$p = 0.740; STIM-vs-PRE d_z = -1.61 |
| 1G | VP$^{GABA}$ HFD consumption — Males [sex-split] | One-way RM-ANOVA + Bonferroni post-hoc | PRE vs STIM vs POST | 5 (5M/0F) | F(2,8) | F = 8.491 | 0.011 | STIM-vs-PRE 0.110 | η$^2$p = 0.680; STIM-vs-PRE d_z = -1.38 |
| 1G | VP$^{GABA}$ HFD consumption — Females [sex-split] | One-way RM-ANOVA + Bonferroni post-hoc | PRE vs STIM vs POST | 7 (0M/7F) | F(2,12) | F = 21.59 | 1.1e-04 | STIM-vs-PRE 0.014 | η$^2$p = 0.783; STIM-vs-PRE d_z = -1.66 |
| 1H | VP$^{GABA}$ chow prestimulation | Paired t-test | laser OFF vs ON (STIM) (pre-access) | 8 (4M/4F) | t(7) | t = -0.321 | 0.758 (ns) | — | Cohen's d_z = -0.11 |
| 1H | VP$^{GABA}$ HFD prestimulation | Paired t-test | laser OFF vs ON (STIM) (pre-access) | 8 (4M/4F) | t(7) | t = 0.351 | 0.736 (ns) | — | Cohen's d_z = 0.12 |
| 1**L** | Arc$^{AgRP}$ photometry AUC | One-way RM-ANOVA (parametric; Shapiro W=0.942, p=0.133) | saline vs ghrelin vs CCK | 9 (5M/4F) | F(2,16) | F = 56.56 | 5.6e-08 | (see post-hoc rows below) | η$^2$p = 0.876 |
| 1**L** | Arc$^{AgRP}$ ghrelin vs saline-baseline | Paired t-test (Holm-corrected) | ghrelin vs saline | 9 (5M/4F) | t(8) | raw p = 2.9e-4 | adj p = 5.8e-4 | — | Cohen's d_z = -2.03 |
| 1**L** | Arc$^{AgRP}$ CCK vs saline-baseline | Paired t-test (Holm-corrected) | CCK vs saline | 9 (5M/4F) | t(8) | raw p = 4.5e-4 | adj p = 5.8e-4 | — | Cohen's d_z = 1.90 |
| 1**L** | Arc$^{AgRP}$ CCK vs ghrelin | Paired t-test (Holm-corrected) | CCK vs ghrelin | 9 (5M/4F) | t(8) | raw p = 1.7e-5 | adj p = 5.2e-5 | — | Cohen's d_z = 3.03 |
| 1**O** | VP$^{GABA}$ photometry AUC | One-way RM-ANOVA (parametric; Shapiro W=0.965, p=0.401) | saline vs ghrelin vs CCK | 10 (3M/7F) | F(2,18) | F = 12.58 | 3.8e-04 | (see post-hoc rows below) | η$^2$p = 0.583 |
| 1**O** | VP$^{GABA}$ ghrelin vs saline-baseline | Paired t-test (Holm-corrected) | ghrelin vs saline | 10 (3M/7F) | t(9) | raw p = 0.124 | adj p = 0.124 (ns) | — | Cohen's d_z = 0.54 |
| 1**O** | VP$^{GABA}$ CCK vs saline-baseline | Paired t-test (Holm-corrected) | CCK vs saline | 10 (3M/7F) | t(9) | raw p = 1.2e-3 | adj p = 3.6e-3 | — | Cohen's d_z = 1.47 |
| 1**O** | VP$^{GABA}$ CCK vs ghrelin | Paired t-test (Holm-corrected) | CCK vs ghrelin | 10 (3M/7F) | t(9) | raw p = 5.8e-3 | adj p = 0.012 | — | Cohen's d_z = 1.14 |
| **Figure 2. VP$^{GABA}$ neurons bi-directionally control reward consumption.** | | | | | | | | | |
| **2C** | Licks per block | Paired t-test | multi-stim activation, laser OFF vs ON | 8 (4M/4F) | t(7) | t = 3.149 | 0.0162 | — | Cohen's d_z = 1.11 |
| **2C** | Bouts per block | Paired t-test | multi-stim activation, laser OFF vs ON | 8 (4M/4F) | t(7) | t = 6.581 | 3e-4 | — | Cohen's d_z = 2.33 |
| **2C** | Average bout duration | Paired t-test | multi-stim activation, laser OFF vs ON | 8 (4M/4F) | t(7) | t = 3.086 | 0.0177 | — | Cohen's d_z = 1.09 |
| **2D** | Licks per block | Paired t-test | continuous activation, laser OFF vs ON | 11 (7M/4F) | t(10) | t = 3.996 | 2.5e-3 | — | Cohen's d_z = 1.21 |
| **2D** | Bouts per block | Paired t-test | continuous activation, laser OFF vs ON | 11 (7M/4F) | t(10) | t = 4.138 | 2e-3 | — | Cohen's d_z = 1.25 |
| **2D** | Average bout duration | Paired t-test | continuous activation, laser | 11 (7M/4F) | t(10) | t = 2.741 | 0.0208 | — | Cohen's d_z = 0.83 |

| | | | | | | | | | |
|---|---|---|---|---|---|---|---|---|---|
| | | | OFF vs ON | | | | | | |
| 2**E** | Licks per block | Paired t-test | constant inhibition (GtACR2), laser OFF vs ON | 9 (6M/3F) | t(8) | t = -3.184 | 0.0129 | — | Cohen's d_z = -1.061 |
| 2**E** | Bouts per block | Paired t-test | constant inhibition (GtACR2), laser OFF vs ON | 9 (6M/3F) | t(8) | t = -2.902 | 0.0198 | — | Cohen's d_z = -0.967 |
| 2**E** | Average bout duration | Paired t-test | constant inhibition (GtACR2), laser OFF vs ON | 9 (6M/3F) | t(8) | t = -3.401 | 9.3e-3 | — | Cohen's d_z = -1.134 |
| 2**F-H** | $VP^{GABA:ChR2}$ single-unit classification | One-sample Wilcoxon vs 0 per unit, BH-FDR (α=0.10), $\lvert$median ΔFR$\rvert$>1 Hz gate | post-stim ΔFR vs 0, per untagged single unit | 10 untagged SU (n=3 mice, 3M/0F) | per-unit (40 bins) | W = 18–387 | per-unit p_FDR 1.7e-5–0.757 | BH-FDR | 0 activated / 5 inhibited / 5 unmodulated; + 3 optically tagged units (defined upstream by light-evoked latency) → 13 single units total |
| **Figure 3. Single-photon endoscopic imaging reveals functionally heterogeneous $VP^{GABA}$ dynamics that encode and track consummatory licking.** | | | | | | | | | |
| 3C | Combined cell classification (lick-bouts 0–3 s) | One-sample Wilcoxon vs 0 per cell, BH-FDR | post-onset z-score vs 0 (339 cells) | 8 (7M/1F); 339 cells | per-cell | median raw p = 5.6e-09 | 297/339 sig (FDR) | — | 129 activated / 168 inhibited / 42 unmodulated |
| 3E | Cell classification, single licks (0–3 s) | One-sample Wilcoxon vs 0 per cell, BH-FDR | post-event z-score vs 0 (339 cells) | 8 (7M/1F); 339 cells | per-cell | median raw p = 5.6e-05 | 252/339 sig (FDR) | — | 110 activated / 142 inhibited / 87 unmodulated |
| 3E | Cell classification, short bouts (0–6 s) | One-sample Wilcoxon vs 0 per cell, BH-FDR | post-event z-score vs 0 (339 cells) | 8 (7M/1F); 339 cells | per-cell | median raw p = 9.0e-08 | 294/339 sig (FDR) | — | 141 activated / 153 inhibited / 45 unmodulated |
| 3E | Cell classification, long bouts (0–15 s) | One-sample Wilcoxon vs 0 per cell, BH-FDR | post-event z-score vs 0 (339 cells) | 8 (7M/1F); 339 cells | per-cell | median raw p = 1.3e-25 | 323/339 sig (FDR) | — | 151 activated / 172 inhibited / 16 unmodulated |
| 3F | Cell response composition | Chi-square test of independence | act/unmod/inh × single/short/long | 8 (7M/1F); 339 cells | $\chi^2(4)$ | $\chi^2$ = 61.44 | 1.4e-12 | — | Cramér's V = 0.174 |
| 3F | Mean activated amplitude | One-way RM-ANOVA (per-mouse) + Holm paired t | single vs short vs long bout | 8 (7M/1F) | F(2,14) | F = 25.34 | 2.2e-5 | S-Sh 0.040; S-L 0.0028; Sh-L 0.0038; s – single, Sh – short, L – long | $\eta^2$p = 0.784 |
| 3F | Mean inhibited amplitude | One-way RM-ANOVA (per-mouse) + Holm paired t | single vs short vs long bout | 8 (7M/1F) | F(2,14) | F = 11.07 | 1.3e-3 | S-Sh 0.313; S-L 0.031; Sh-L 0.023 | $\eta^2$p = 0.613 |
| 3H | Licks per block | Paired t-test | closed-loop, laser OFF vs ON | 15 (9M/6F) | t(14) | t = 3.659 | 2.6e-3 | — | Cohen's d_z = 0.945 |
| 3**I** | Bouts per block | Paired t-test | closed-loop, laser OFF vs ON | 15 (9M/6F) | t(14) | t = -1.674 | 0.1164 (ns) | — | Cohen's d_z = 0.432 |
| 3**J** | Average bout duration | Paired t-test | closed-loop, laser OFF vs ON | 15 (9M/6F) | t(14) | t = 6.632 | <0.0001 | — | Cohen's d_z = 1.712 |
| **Figure 4. $VP^{GABA}$ neurons are required for the development of diet-induced obesity.** | | | | | | | | | |
| 4B | slc17a6 (VGLUT2) transcript count | Independent t-test | mCherry vs taCasp3 | mCh 5 (2M/3F); casp 5 (2M/3F) | t(8) | t = -0.321 | 0.757 (ns) | — | Cohen's d = -0.20 |
| 4B | slc32a1 (VGAT/GABA) transcript count | Independent t-test | mCherry vs taCasp3 | mCh 5 (2M/3F); casp 5 (2M/3F) | t(8) | t = 4.721 | 1.5e-03 | — | Cohen's d = 2.99 |
| 4C | Cumulative licks at 15 min‡ | Independent t-test | mCherry vs taCasp3 | mCh 11 (5M/6F); casp 13 (8M/5F) | t(22) | t = 2.885 | 8.6e-03 | — | Cohen's d = 1.18 |
| 4E | Avg lick bouts/session | Independent t-test | mCherry vs taCasp3 | mCh 11 (5M/6F); casp 13 (8M/5F) | t(22) | t = 2.636 | 0.015 | — | Cohen's d = 1.08 |
| 4E | Mean lick-bout duration | Independent t-test | mCherry vs taCasp3 | mCh 11 (5M/6F); casp 13 (8M/5F) | t(22) | t = 0.870 | 0.393 (ns) | — | Cohen's d = 0.36 |
| 4F | Per-mouse modal lick frequency | Independent t-test | mCherry vs taCasp3 | mCh 11 (5M/6F); casp 13 (8M/5F) | t(22) | t = 3.091 | 5.3e-03 | — | Cohen's d = 1.27 |
| 4G | Diet consumption Group effect | Mixed RM-ANOVA (between Group, within State) | mCherry vs taCasp3 | 18 (10 mCh / 8 casp) | F(1,16) | F = 1.789 | 0.200 (ns) | — | $\eta^2$p = 0.101 |
| 4G | Diet consumption State effect | Mixed RM-ANOVA | Sated vs Fasted | 18 | F(1,16) | F = 59.53 | 8.8e-07 | — | $\eta^2$p = 0.788 |
| 4G | Diet consumption Group × State | Mixed RM-ANOVA | Group × State interaction | 18 | F(1,16) | F = 6.694 | 0.020 | — | $\eta^2$p = 0.295 |
| 4G | Consumption post-hoc (Sated) | Mann-Whitney U, Holm-corrected | mCherry vs taCasp3 (Sated) | mCh 8 / casp 10 | — | U = 67.0 | raw 0.0185 | p_Holm = 0.037 | rank-biserial r = -0.68 |
| 4G | Consumption post-hoc (Fasted) | Mann-Whitney U, Holm-corrected | mCherry vs taCasp3 (Fasted) | mCh 8 / casp 10 | — | U = 41.5 | raw 0.929 | p_Holm = 0.929 (ns) | rank-biserial r = -0.04 |
| 4I | HFD weight Wk1→Wk5, mCherry-HFD | Paired t-test | week 1 vs week 5 | 23 (15M/8F) | t(22) | t = -4.538 | 1.6e-04 | — | Cohen's d_z = -0.95 |
| 4I | HFD weight Wk1→Wk5, taCasp3-HFD | Paired t-test | week 1 vs week 5 | 22 (17M/5F) | t(21) | t = 1.269 | 0.218 (ns) | — | Cohen's d_z = 0.27 |
| 4I | HFD weight Wk1→Wk5, taCasp3-chow | Paired t-test | week 1 vs week 5 | 7 (2M/5F) | t(6) | t = -1.747 | 0.131 (ns) | — | Cohen's d_z = -0.66 |
| 4J | HFD weight Wk5→Wk10, mCherry-HFD | Paired t-test | week 5 vs week 10 | 23 (15M/8F) | t(22) | t = -8.752 | 1.1e-08 | — | Cohen's d_z = -1.83 |
| 4J | HFD weight Wk5→Wk10, taCasp3-HFD | Paired t-test | week 5 vs week 10 | 22 (17M/5F) | t(21) | t = -6.531 | 1.7e-06 | — | Cohen's d_z = -1.39 |
| 4J | HFD weight Wk5→Wk10, taCasp3-chow | Paired t-test | week 5 vs week 10 | 7 (2M/5F) | t(6) | t = -2.969 | 0.025 | — | Cohen's d_z = -1.12 |
| 4J | HFD weight Wk10 group comparison | Mann-Whitney U | mCherry-HFD vs taCasp3-HFD at week 10 | mCh-HFD 23 (15M/8F) / casp-HFD 22 (17M/5F) | — | U = 427.0 | 8.2e-05 | — | rank-biserial r = -0.69 |
| **4J** | HFD weight Wk10 group | Mann-Whitney U | mCherry-HFD vs taCasp3- | mCh-HFD (15M) / | — | U = 221.0 | <0.001 | — | rank-biserial r = -0.733 |

| | | | | | | | | | |
|---|---|---|---|---|---|---|---|---|---|
| | comparison (sex split, males) | | HFD at week 10 | casp-HFD (17M) | | | | | |
| **4J** | HFD weight Wk10 group comparison (sex split, females) | Mann-Whitney U | mCherry-HFD vs taCasp3-HFD at week 10 | mCh-HFD (8F) / casp-HFD (5F) | — | U = 39.0 | 0.003 | — | rank-biserial r = -0.950 |
| **Supplemental Figure 1. Behavioral ethograms and locomotor tracking during optogenetic VP$^{GABA}$ stimulation.** | | | | | | | | | |
| S1B | Chow interaction duration | One-way RM-ANOVA + Holm post-hoc | PRE vs STIM vs POST | 10 (6M/4F) | F(2,18) | F = 6.254 | 0.0087 | PRE-STIM 0.095<br>STIM-POST 0.095<br>PRE-POST 0.505 | $\eta^2$p = 0.410; STIM-vs-PRE d_z = -0.80 |
| S1B | HFD interaction duration | One-way RM-ANOVA + Holm post-hoc | PRE vs STIM vs POST | 10 (6M/4F) | F(2,18) | F = 9.199 | 0.0018 | PRE-STIM 0.042<br>STIM-POST 0.042<br>PRE-POST 0.933 | $\eta^2$p = 0.506; STIM-vs-PRE d_z = -0.96 |
| S1B | Non-food object interaction duration | One-way RM-ANOVA + Holm post-hoc | PRE vs STIM vs POST | 9 (5M/4F) | F(2,16) | F = 4.046 | 0.0378 | PRE-ON 0.231<br>ON-POST 0.231<br>PRE-POST 0.924 | $\eta^2$p = 0.336; ON-vs-PRE d_z = -0.67 |
| S1D | Average locomotor speed | Paired t-test | laser OFF vs ON | 8 (4M/4F) | t(7) | t = -0.872 | 0.412 (ns) | — | Cohen's d_z = -0.31 |
| S1D | Time in arena centre | Paired t-test | laser OFF vs ON | 8 (4M/4F) | t(7) | t = -2.462 | 0.043 | — | Cohen's d_z = -0.87 |
| **Supplemental Figure 2. Single cell RNA-sequencing of the VP and whole-brain mapping of fasting-activated inputs to the VP.** | | | | | | | | | |
| S2A | Feeding gene enrichment between Arc and VP | AddModuleScore (Seurat); n=100 control gene randomly selected | Arc vs. VP expression | — | — | — | — | — | — |
| S2D-F | Whole brain cFos counts | Light sheet imaging + cFos nuclei counting | See table S2 | 8 (8M/0F), per group | See table S2 | See table S2 | See table S2 | See table S2 | See table S2 |
| **Supplemental Figure 3. Fiber photometry of Arc$^{AgRP}$ and VP$^{GABA}$ neurons aligned to pellet retrievals dispensed from FED3.** | | | | | | | | | |
| S3B | Post-injection pellet intake | Paired t-test | Saline vs ghrelin | 9 (7M/2F) | t(8) | t = 12.48 | 1.6e-6 | — | Cohen's d_z = 4.16 |
| S3E | Peak deflection (-1s to +1s) | Welch's t-test | Arc$^{AgRP}$ vs VP$^{GABA}$ (between-subjects) | Arc$^{AgRP}$ (9, 5M/4F); VP$^{GABA}$ (6, 4M/2F) | df = 11.43 | t = -18.394 | <0.0001 | — | d = -8.982 |
| S3F | Post-event AUC (Arc$^{AgRP}$) | Paired t-test | First vs last retrieval | 9 (5M/4F) | t(8) | t = -3.851 | 4.9e-03 | — | Cohen's d_z = -1.284 |
| S3G | Post-event AUC (VP$^{GABA}$) | Paired t-test | First vs last retrieval | 6 (4M/2F) | t(5) | t = 1.837 | 0.1257 | — | Cohen's d_z = 0.750 |
| **Supplemental Figure 4. VP$^{GABA}$ optical activation and inhibition drive preference and avoidance, respectively.** | | | | | | | | | |
| S4C | Time on stim-paired side (%), ChR2 | Paired t-test | baseline (OFF) vs laser-ON | 13 (8M/5F) | t(12) | t = -7.068 | 1.3e-05 | — | Cohen's d_z = -1.96 |
| S4F | Time on stim-paired side (%), GtACR2 | Paired t-test | baseline (OFF) vs laser-ON | 8 (5M/3F) | t(7) | t = 3.974 | 5.4e-03 | — | Cohen's d_z = 1.41 |
| **Supplemental Figure 5. Optogenetic activation of VP$^{GABA}$ neurons drives licking for quinine and an empty spout.** | | | | | | | | | |
| S5B | Quinine licks per block | Paired t-test | laser OFF vs ON | 8 (4M/4F) | t(7) | t = 5.761 | 6.9e-04 | — | Cohen's d_z = 2.04 |
| S5B | Quinine bouts per block | Paired t-test | laser OFF vs ON | 8 (4M/4F) | t(7) | t = 4.732 | 2.1e-03 | — | Cohen's d_z = 1.67 |
| S5B | Quinine average bout duration | Paired t-test | laser OFF vs ON | 8 (4M/4F) | t(7) | t = 5.008 | 1.6e-03 | — | Cohen's d_z = 1.77 |
| S5D | Empty licks per block | Paired t-test | laser OFF vs ON | 10 (5M/5F) | t(9) | t = 3.324 | 8.9e-03 | — | Cohen's d_z = 1.05 |
| S5D | Empty bouts per block | Paired t-test | laser OFF vs ON | 10 (5M/5F) | t(9) | t = 4.267 | 2.1e-03 | — | Cohen's d_z = 1.35 |
| S5D | Empty average bout duration | Paired t-test | laser OFF vs ON | 10 (5M/5F) | t(9) | t = 3.329 | 8.8e-03 | — | Cohen's d_z = 1.05 |
| S5F | Cross-condition licks per block | One-way ANOVA + Welch post-hoc (uncorrected) | quinine vs empty vs boost (laser-ON) | 29 (16M/13F) | F(2,26) | F = 4.855 | 0.016 | q-empty 0.138<br>q-boost 0.0077<br>empty-boost 0.0775 | $\eta^2$p = 0.272 |
| S5F | Cross-condition bouts per block | One-way ANOVA | quinine vs empty vs boost | 29 (16M/13F) | F(2,26) | F = 0.494 | 0.616 (ns) | — | $\eta^2$p = 0.037 |
| S5F | Cross-condition average bout duration | One-way ANOVA + Welch post-hoc (uncorrected) | quinine vs empty vs boost | 29 (16M/13F) | F(2,26) | F = 2.984 | 0.068 (ns) | q-empty 0.071; q-boost 0.039; empty-boost 0.164 | $\eta^2$p = 0.187 |
| S5G | Cross-condition modal lick freq | One-way ANOVA | quinine vs empty vs boost | 29 (16M/13F) | F(2,26) | F = 0.981 | 0.388 (ns) | — | $\eta^2$p = 0.070 |
| **Supplemental Figure 6. Principal component state dynamics of VP$^{GABA}$ neurons and linear decoding of VP$^{GABA}$ population dynamics during self-paced drinking.** | | | | | | | | | |
| S6C | Coefficients of PC1 eigenvector | Top and bottom quartile (25% hi/lo) | — | — | — | — | — | — | — |
| S6D | High PC1 (top 25%) post-event AUC | Paired t-test (Holm) | Single vs short | 84 cells (8 mice, 7M/1F) | t(83) | t = -6.006 | < 0.0001 | < 0.0001 | Cohen's d_z = -0.655 |
| S6D | High PC1 (top 25%) post-event AUC | Paired t-test (Holm) | Single vs long | 84 cells (8 mice, 7M/1F) | t(83) | t = -8.100 | < 0.0001 | < 0.0001 | Cohen's d_z = -0.884 |
| S6D | High PC1 (top 25%) post-event AUC | Paired t-test (Holm) | Short vs long | 84 cells (8 mice, 7M/1F) | t(83) | t = -5.596 | < 0.0001 | < 0.0001 | Cohen's d_z = -0.611 |
| S6D | Low PC1 (bottom 25%) post-event AUC | Paired t-test (Holm) | Single vs short | 84 cells (8 mice, 7M/1F) | t(83) | t = 7.943 | < 0.0001 | < 0.0001 | Cohen's d_z = 0.867 |
| S6D | Low PC1 (bottom 25%) post-event AUC | Paired t-test (Holm) | Single vs long | 84 cells (8 mice, 7M/1F) | t(83) | t = 17.027 | < 0.0001 | < 0.0001 | Cohen's d_z = 1.858 |
| S6D | Low PC1 (bottom 25%) post-event AUC | Paired t-test (Holm) | Short vs long | 84 cells (8 mice, 7M/1F) | t(83) | t = 13.370 | < 0.0001 | < 0.0001 | Cohen's d_z = 1.459 |
| S6G, H | Balanced accuracy vs shuffle (time resolved) | Wilcoxon signed rank per dt (BH-FDR) | Decoder > shuffle | 8 (7M/1F); 13 sessions | — | — | — | Min FDR p = 0.0004 | — |
| S6I | auROC, -3 to +0.5 s window | One-sample Wilcoxon vs 0.5 (BH-FDR) | per-session auROC vs chance | 8 (7M/1F); 13 sessions | n=13 | median auROC = 0.490 | p_raw 0.787; FDR 0.893 (ns) | — | median 0.490 vs 0.5 |
| S6I | auROC, -3 to +1 s window | One-sample Wilcoxon vs 0.5 | per-session auROC vs | 8 (7M/1F); 13 | n=13 | median auROC | p_raw 0.893; | — | median 0.490 vs 0.5 |

| | | | | | | | | | |
|---|---|---|---|---|---|---|---|---|---|
| | | (BH-FDR) | chance | sessions | | = 0.490 | FDR 0.893 (ns) | | |
| S6I | auROC, -3 to +3 s window | One-sample Wilcoxon vs 0.5 (BH-FDR) | per-session auROC vs chance | 8 (7M/1F); 13 sessions | n=13 | median auROC = 0.595 | p_raw 0.216; FDR 0.433 (ns) | — | median 0.595 vs 0.5 |
| S6I | auROC, -3 to +10 s window | One-sample Wilcoxon vs 0.5 (BH-FDR) | per-session auROC vs chance | 8 (7M/1F); 13 sessions | n=13 | median auROC = 0.731 | p_raw 1.2e-3; FDR 4.9e-3 | — | median 0.731 vs 0.5 |
| **Supplemental Figure 7. VP$^{GABA}$ ablation leaves homeostatic weight regulation intact, but HFD exposure in VP$^{GABA}$ ablated mice induces hyperglycemia.** | | | | | | | | | |
| S7B | Chow weight Wk1→Wk5, mCherry-chow | Paired t-test | week 1 vs week 5 | 5 (0M/5F) | t(4) | t = -0.787 | 0.475 (ns) | — | Cohen's d_z = -0.35 |
| S7B | Chow weight Wk1→Wk5, taCasp3-chow | Paired t-test | week 1 vs week 5 | 7 (2M/5F) | t(6) | t = -1.747 | 0.131 (ns) | — | Cohen's d_z = -0.66 |
| S7C | Chow weight Wk5→Wk10, mCherry-chow | Paired t-test | week 5 vs week 10 | 5 (0M/5F) | t(4) | t = -3.633 | 0.022 | — | Cohen's d_z = -1.63 |
| S7C | Chow weight Wk5→Wk10, taCasp3-chow | Paired t-test | week 5 vs week 10 | 7 (2M/5F) | t(6) | t = -2.969 | 0.025 | — | Cohen's d_z = -1.12 |
| S7C | Chow weight Wk10 group comparison | Mann-Whitney U | control-chow vs taCasp3-chow at week 10 | ctrl-chow 5 / casp-chow 7 | — | U = 27.0 | 0.149 (ns) | — | rank-biserial r = -0.54 |
| S7F | Average daily pellets | Independent t-test (two-tailed) | mCherry vs taCasp3 | mCh 5 (0M/5F); casp 7 (2M/5F) | t(10) | t = -1.184 | 0.264 (2-tailed); 0.868 (1-tailed) | — | Cohen's d = -0.69 |
| S7H | GTT area under the curve | One-way ANOVA + Tukey HSD | chow (mCherry, taCasp3 combined) vs mCherry-HFD vs taCasp3-HFD | 43: chow 12 (2M/10F); mCh-HFD 14 (8M/6F); casp-HFD 17 (10M/7F) | F(2,40) | F = 30.94 | 7.6e-09 | chow vs mCh-HFD <0.001<br>chow vs casp-HFD <0.001<br>mCh-HFD vs casp-HFD 0.0173 | η²p = 0.607 |
| **Supplemental Figure 8. VP$^{GABA}$ ablation does not disrupt circadian feeding patterns.** | | | | | | | | | |
| S8A | Cumulative hopper touches (72 h) | Independent t-test (two-tailed) | mCherry vs taCasp3 | 31 (18M/13F): mCh 16 (8/8); casp 15 (10/5) | t(29) | t = -0.498 | 0.622 (ns) | — | Cohen's d = -0.18 |
| S8D | Daily kcal consumed | Independent t-test (one-tailed) | control vs taCasp3 | 42 (31M/11F): ctrl 21 (14/7); casp 21 (17/4) | t(40) | t = 1.881 | 0.034 (1-tailed); 0.067 (2-tailed) | — | Cohen's d = 0.58 |
| S8E | Daily meal counts | Independent t-test (one-tailed) | mCherry vs taCasp3 | 31 (18M/13F): mCh 16; casp 15 | t(29) | t = -0.389 | 0.700 (2-tailed) | — | Cohen's d = -0.14 |
| S8F | Average meal duration | Independent t-test (one-tailed) | mCherry vs taCasp3 | 31 (18M/13F): mCh 16; casp 15 | t(29) | t = 0.817 | 0.420 (2-tailed) | — | Cohen's d = 0.29 |
| **Supplemental Figure 9. Effect of VP$^{GABA}$ ablation on operant reward seeking and reversal learning.** | | | | | | | | | |
| S9C | Pokes per pellet (animal-averaged) | Independent t-test | taCasp3 vs mCherry | 12 (taCasp3 7 / mCh 5) | t(10) | t = 0.538 | 0.602 (ns) | — | Cohen's d = 0.32 |
| S9C | Median break point (animal-averaged) | Independent t-test | taCasp3 vs mCherry | 12 (taCasp3 7 / mCh 5) | t(10) | t = 0.610 | 0.556 (ns) | — | Cohen's d = 0.36 |
| S9G | Pre-reversal accuracy | Independent t-test | mCherry vs taCasp3 | 12 (5/7) | t(10) | t = 2.278 | 0.046 | — | Cohen's d = 1.33 |
| S9H | Pellets/day | Independent t-test (two-sided) | mCherry vs taCasp3 | 12 (5/7) | t(10) | t = 0.097 | 0.924 (ns) | — | Cohen's d = 0.06 |
| S9I | Logistic-regression OR, lag t-5 | Independent t-test | mCherry vs taCasp3 | 12 (5/7) | t(10) | t = 0.760 | 0.465 (ns) | — | Cohen's d = 0.45 |
| S9I | Logistic-regression OR, lag t-4 | Independent t-test | mCherry vs taCasp3 | 12 (5/7) | t(10) | t = 1.746 | 0.111 (ns) | — | Cohen's d = 1.02 |
| S9I | Logistic-regression OR, lag t-3 | Independent t-test | mCherry vs taCasp3 | 12 (5/7) | t(10) | t = -0.605 | 0.559 (ns) | — | Cohen's d = -0.35 |
| S9I | Logistic-regression OR, lag t-2 | Independent t-test | mCherry vs taCasp3 | 12 (5/7) | t(10) | t = 0.995 | 0.343 (ns) | — | Cohen's d = 0.58 |
| S9I | Logistic-regression OR, lag t-1 | Independent t-test | mCherry vs taCasp3 | 12 (5/7) | t(10) | t = 1.844 | 0.095 (ns) | — | Cohen's d = 1.08 |
| S9J | Win-stay | Independent t-test | mCherry vs taCasp3 | 12 (5/7) | t(10) | t = 1.846 | 0.095 (ns) | — | Cohen's d = 1.08 |
| S9K | Lose-stay | Independent t-test | mCherry vs taCasp3 | 12 (5/7) | t(10) | t = 0.668 | 0.520 (ns) | — | Cohen's d = 0.39 |
| **Supplemental Figure 10. Social seeking behaviors remain intact following VP$^{GABA}$ ablation.** | | | | | | | | | |
| S10B | Average social seeking event (Day 2-4 averaged per animal) | Mann-Whitney U | mCherry vs. taCasp3 | 5 mCh (0M/5F); 7 casp (2M/5F) | — | U = 20 | 0.755 (ns) | — | Rank-biserial r = 0.143 |
| S10D | Social seeking AUC during the dark cycle (Day 2-4 averaged per animal) | Mann-Whitney U | mCherry vs. taCasp3 | 5 mCh (0M/5F); 7 casp (2M/5F) | — | U = 22 | 0.530 (ns) | — | Rank-biserial r = -0.257 |
| S10F | Percentage of approach trials after nose poke (Day 2-4) | Mann-Whitney U | mCherry vs. taCasp3 | 5 mCh (0M/5F); 7 casp (2M/5F) | — | U = 15 | 0.755 (ns) | — | Rank-biserial r = -0.143 |
| S10G | Median approach latency after nose poke (Day 2-4) | Mann-Whitney U | mCherry vs. taCasp3 | 5 mCh (0M/5F); 7 casp (2M/5F) | — | U = 30 | 0.048 | — | Rank-biserial r – 0.714 |
| S10D | Home-cage activity, lights OFF (Night) | Mann-Whitney U, Holm-corrected | mCherry vs taCasp3 | 45 (32M/13F): mCh 23; casp 22 | — | U = 116.0 | raw 0.0019; Holm 0.0039 | — | rank-biserial r = 0.54; d = -1.11 |
